# Selective Prespacer Processing Ensures Precise CRISPR-Cas Adaptation

**DOI:** 10.1101/608976

**Authors:** Sungchul Kim, Luuk Loeff, Sabina Colombo, Stan J.J. Brouns, Chirlmin Joo

## Abstract

CRISPR-Cas immunity protects prokaryotes against foreign genetic elements. CRISPR-Cas uses the highly conserved Cas1-Cas2 complex to establish inheritable memory (spacers). It remains elusive how Cas1-Cas2 acquires spacers from cellular DNA fragments (prespacers) and how it integrates them into the CRISPR array in the correct orientation. By using the high spatiotemporal resolution of single-molecule fluorescence, we reveal that Cas1-Cas2 obtains prespacers in various forms including single-stranded DNA and partial duplexes by selecting them in the DNA-length and PAM-dependent manner. Furthermore, we identify DnaQ exonucleases as enzymes that can mature the Cas1-Cas2-loaded precursor prespacers into an integration-competent size. Cas1-Cas2 protects the PAM sequence from maturation, which results in the production of asymmetrically trimmed prespacers and subsequent spacer integration in the correct orientation. This kinetic coordination in prespacer selection and PAM trimming provides comprehensive understanding of the mechanisms that underlie the integration of functional spacers in the CRISPR array.

## Introduction

CRISPR (clustered regularly interspaced short palindromic repeats)-Cas (CRISPR-associated) systems are RNA-guided adaptive immune systems that defend prokaryotes against invading nucleic acids (Hille et al., 2018; Marraffini, 2015). To fulfil inheritable immunity against invaders, CRISPR-Cas systems undergo three central steps. The first step (adaptation or spacer acquisition) is characterized by an update of the genetic memory. In this step, foreign DNA fragments (prespacers) are recognized and integrated between the repeats of the CRISPR array to form spacers (Barrangou et al., 2007; Fineran and Charpentier, 2012; Jackson et al., 2017; McGinn and Marraffini, 2019). In the second step (expression), the spacers are transcribed into long precursor CRISPR RNAs (pre-crRNAs) and processed into CRISPR RNAs (crRNAs) that assemble with Cas proteins into ribonucleoprotein complexes such as Cascade, and Cas9 (Marraffini and Sontheimer, 2008; van der Oost et al., 2014). In the final step of CRISPR immunity (interference), invading nucleic acids are bound by the RNA guided surveillance complexes (R-loop formation), resulting in target cleavage that eliminates the invading threat (Deveau et al., 2010; Garneau et al., 2010; Hale et al., 2009; Hille et al., 2018; Marraffini and Sontheimer, 2008). To date, the expression and interference stages of CRISPR–Cas systems have been widely studied, proving detailed insights into the molecular mechanisms underlie these stages. However, the molecular details of how Cas1-Cas2 selects substrates for the integration of spacers remains poorly understood.

Adaptation relies on the highly conserved Cas1 and Cas2 proteins, that form a heterohexameric Cas1_(4)_-Cas2_(2)_ complex (here after called Cas1-Cas2) (Jackson et al., 2017; McGinn and Marraffini, 2019; Nunez et al., 2016; Nunez et al., 2015a; Nunez et al., 2014; Nunez et al., 2015b; Wang et al., 2015; Wright et al., 2016; Xiao et al., 2017). In vivo studies on spacer acquisition have suggested that Cas1-Cas2 identifies suitable prespacers (PS) based on the presence protospacer adjacent motif (PAM), which is also a prerequisite for the CRISPR-interference stage of immunity (Diez-Villasenor et al., 2013; Levy et al., 2015; Yosef et al., 2012). The absence of PAMs in the spacer flanking repeat sequences prevents self-recognition and thereby inhibits autoimmunity.

The sources of PS DNA have been suggested to be diverse. For example, genome wide profiling of naïve adaptation has shown that Cas1-Cas2 derives PS DNA from RecBCD degradation intermediates that are generated during the repair of double-stranded DNA (dsDNA) breaks (Babu et al., 2011; Cubbon et al., 2018; Ivancic-Bace et al., 2015; Levy et al., 2015; Radovcic et al., 2018). Other studies have shown that the degradation products generated by the Cascade-Cas3 complex during interference, can be repurposed by Cas1-Cas2 to facilitate rapid acquisition of new spacers (so called interference-driven primed adaptation) (Datsenko et al., 2012; Fineran et al., 2014; Jackson et al., 2017; Kunne et al., 2016; Musharova et al., 2017; Semenova et al., 2016). Intriguingly, both RecBCD and Cas3 are known to generate single-stranded DNA (ssDNA) degradation products, which contrast with the optimal substrate for integration by Cas1-Cas2 (Kunne et al., 2016; Mulepati and Bailey, 2013; Yeeles et al., 2011a; Yeeles et al., 2011b). For example, the optimal substrate for Escherichia coli (E. coli) Cas1-Cas2 of the type I-E system is composed of a central 23-bp duplex with two 5-nt 3’-overhangs (here after called canonical PS) (Nunez et al., 2015a; Nunez et al., 2014; Wang et al., 2015). This suggests that the fragments generated by RecBCD and Cascade-Cas3 must re-anneal before integration into the CRISPR-array by Cas1-Cas2. It will be of great interest to find out whether Cas1-Cas2 may play an active role in capturing the ssDNA fragments from these sources to form an effective Cas1-Cas2-PS integration complex.

Structural studies of E. coli type I-E Cas1-Cas2 bound to a PS have shown that the PAM sequence (5’-CTT-3’), which is required for PS selection, must be located 5-nt away from the duplex end (at nucleotide +5 to +7 in the 3’-overhang) (Wang et al., 2015). This suggests that in order to recognize the PAM sequence, Cas1-Cas2 should bind PS precursors with 3’-overhangs that are longer or equal to 7-nt. As a consequence, 3’-overhang of the precursor PS must be processed into the optimal size of 5-nt with a 3’-hydroxyl group for integration into the CRISPR array (Jackson et al., 2017; Nunez et al., 2015b; Wang et al., 2015). To date, it remains unclear how Cas1-Cas2 can recognize and load PS precursors containing a PAM. Moreover, it is also unknown how the PAM-containing 3’-overhangs of PS precursors are matured for PAM-specific spacer integration.

In the final stage of spacer acquisition, spacers must be integrated in the correct orientation with respect to position of the PAM sequence in the 3’-overhang (Nunez et al., 2015a; Nunez et al., 2015b). In vitro, Cas1-Cas2 has been shown to integrate spacers in either orientation with an equal probability (Lopez-Sanchez et al., 2012; Nunez et al., 2016; Shmakov et al., 2014; Xiao et al., 2017). However, in vivo only correctly oriented spacers with respect to PAM sequence will result in functional crRNAs for target recognition (Jackson et al., 2017; McGinn and Marraffini, 2019). Recent in vivo and in vitro studies using type I-A, I-B, I-C and I-D revealed the critical role of Cas4 in PS DNA maturation and the fidelity of spacer integration (Hou and Zhang, 2018; Kieper et al., 2018; Lee et al., 2018; Rollie et al., 2018; Shiimori et al., 2018). However, it is unknown how Cas4-deficient systems such as E. coli type I-E determine the correct orientation of new spacers.

Here, we employed single-molecule and biochemical assays to illuminate the entire process of spacer acquisition—from PS precursor selection to 3’-overhang processing and subsequent integration of new spacers into the CRISPR array. We show that Cas1-Cas2 kinetically selects PAM-containing precursor DNAs from various sources of DNA, including ssDNA and partially duplexed DNA with long flanking 3’-overhangs. Moreover, we identify DNA polymerase III (Pol III) and Exonuclease T (also known as ExoT or RNase T) as trimming enzymes that are capable of maturing the ssDNA flanks of precursor DNA with extended 3’-overhangs. The recognition of the PAM sequence by Cas1-Cas2 is mediated by the C-terminal tail of Cas1, which determines when the 3’-overhang is trimmed. This asymmetry in 3’-overhang trimming ensures that the integration of the PAM-derived 3’-end of a PS occurs after that of a non-PAM end. This PAM-dependent two-step process results in the effective spacer integration in the correct orientation.

## Results

### Single-Molecule Observation of DNA Loading by Cas1-Cas2

The Cas1-Cas2 integration complex repurposes DNA fragments from invading genetic elements (Kunne et al., 2016; Levy et al., 2015). The majority of these DNA fragments will have structures that deviate from the canonical form of a PS DNA: a central 23-bp duplex with two 5-nt 3’-overhangs. Therefore, the Cas1-Cas2 integration complex should be able to bind non-canonical PS DNAs, hereafter called precursor PS DNAs, as a first step towards integration into the CRISPR array. To elucidate the mechanism for PS precursors selection by the Cas1-Cas2 integration complex, we sought to visualize precursor PS loading by Cas1-Cas2 with a high spatiotemporal resolution.

To image PS precursor loading by Cas1-Cas2 complexes, we developed an assay based on single-molecule Förster resonance energy transfer (smFRET). In brief, Cas1-Cas2 complexes were site-specifically labelled with a biotin at the N-terminus of Cas1 and immobilized to a polyethylene glycol (PEG)-coated quartz slide through biotin-streptavidin linkage (Figures 1A and S1A). The immobilized Cas1-Cas2 complexes were presented with DNA substrates that were labelled with a donor (Cy3) and an acceptor (Cy5) dye on the top and bottom strands, respectively (Figure 1B and Table S1). This scheme yielded a FRET value of ~0.72 (Figure S1B) and enabled us to probe binding events in real time by collecting the fluorescence signals through total internal reflection fluorescence (TIRF) microscopy (Figure 1A).

**Figure 1:**
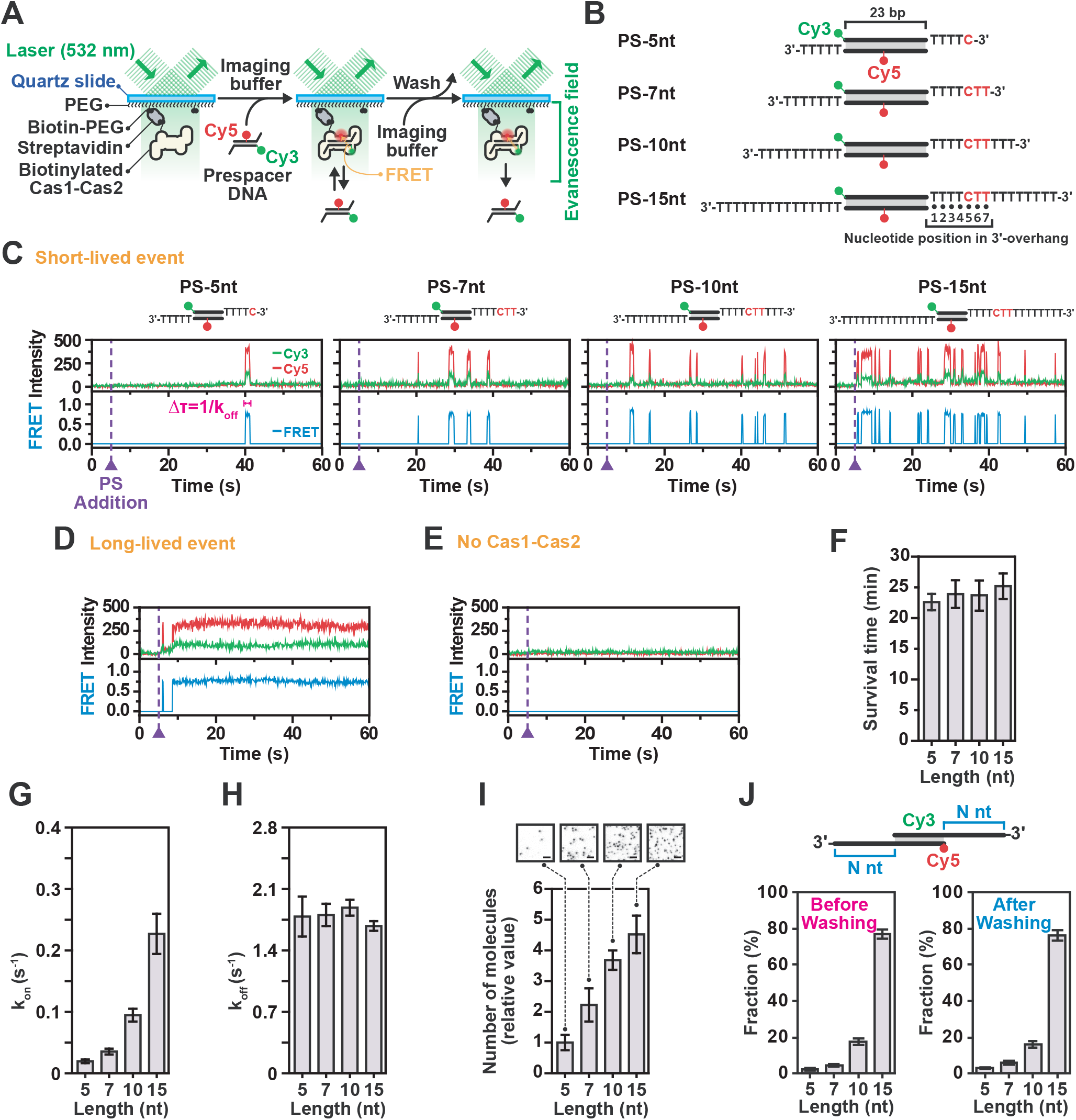
3’-Overhang Length-Dependent Prespacer Selection by Cas1-Cas2. **(A)** Schematic design of single-molecule assay used to monitor binding of Cas1-Cas2 to PS DNAs. **(B)** Design of PS DNA constructs used for 3’-overhang length-dependent single-molecule assays. These DNA constructs commonly consists of 23-bp central duplex, Cy3-labelled top PS strands at 5’-ends and bottom strands Cy5-labelled at 16th T nucleotide from the 5’-ends. **(C)** Representative time traces of donor (Cy3, green) and acceptor (Cy5, red) fluorescence and corresponding FRET (blue) exhibiting short-lived PS DNA binding events for various 3’-overhang lengths at a single spot in the microfluidic chamber. The duration of each binding event was measured as the dwell time (Δτ). DNA was added at time = 5 s. **(D)** Representative time trace of long-lived PS DNA binding. DNA was added at time = 5 s. **(E)** Representative time traces in absence of Cas1-Cas2. DNA was added at time = 5 s. **(F)** Survival time of stably bound PS DNA on Cas1-Cas2 over various 3’-overhang lengths washed after 30 min of incubation. Data represents mean ± SEM (n = 3). **(G)** The k_on_ values were calculated from cumulative probability of arrival time over various PS 3’-overhang lengths (right). Error bars indicate the 95% confidence interval obtained by bootstrap analysis. **(H)** Dissociation rate (k_off_) of short-lived bindings over various PS 3’-overhang lengths. The koff values were calculated from dwell time (Δτ) of each event by single-exponential decay fitting. Error bars indicate the 95% confidence interval obtained by bootstrap analysis. **(I)** Quantification of the number of molecules over various 3’-overhang lengths at 30 min after PS DNA addition. Data represents mean ± SEM (n = 3). Representative CCD images (acceptor channel) are included as in-sets. **(J)** Fractions of each FRET population of Cas1-Cas2-bound PS molecules various 3’-overhang lengths at 30 min after PS DNA addition and followed by washing to observe stably bound molecules. Data represents mean ± SEM (n = 3).

We probed the binding properties of Cas1-Cas2 to a canonical PS DNAs with a 5-nt 3’-overhang (PS-5nt) (Figure 1B) that is primed for spacer integration into to the CRISPR array (Nunez et al., 2015a; Nunez et al., 2014; Nunez et al., 2015b; Wang et al., 2015). Unexpectedly, the majority of binding events (>99%) showed transient interactions that were marked by sudden appearance and subsequent rapid disappearance of the fluorescence intensity (Figure 1C), whereas long stable binding events were rarely observed (<1%) (Figure 1D). Notably, control experiments in the absence of the Cas1-Cas2 complex showed no appreciable binding of the PS DNA (Figure 1E). Analysis of the transient interactions resulted in a histogram that followed a single-exponential decay with a characteristic dwell time (Δτ) of 0.560 ± 0.072 s (Figure S1C). In contrast, the long-lived population persisted over the entire imaging duration (~1 minute) or until photobleached (Figure 1D). Survival rate analysis of this long-lived population displayed a single-exponential decay with a characteristic dwell-time (Δτ) of 22.5 ± 1.3 minutes (Figures 1F and S1D). These results suggest that the Cas1-Cas2 complex requires frequent interactions with a DNA substrate before transitioning to a stably bound state.

### Cas1-Cas2 Selects Prespacers with Long 3’-Overhangs

Next, we probed the properties of Cas1-Cas2 binding to a series of precursor PS DNAs with 3’-overhangs in increasing lengths (7-nt, 10-nt and 15-nt) (Figure 1B). Our single-molecule assay revealed that the length of the 3’-overhang dictates the binding frequency of Cas1-Cas2, resulting in more frequent binding as the length of the 3’-overhang increased (Figure S1E). Kinetic analysis of these binding events showed that the binding frequency (k_on_) exponentially increased with a longer 3’-overhang (Figure 1G), whereas the dwell-time (1/k_off_) remained unaltered (Figures 1H and S1C). As a result of the more frequent interactions with the precursor PS DNA, the number of stably bound molecules increased proportional with the length of the 3’-overhang (Figure 1I), while the survival rate remained constant (Figures 1F and S1D). These results are consistent with the binding behavior in electromobility shift assays (EMSA) (Figure S1F) and previously published studies (Moch et al., 2017; Wang et al., 2015) and further suggests that Cas1-Cas2 kinetically selects for PS precursors with long 3’-overhangs through frequent interactions.

In the cell, DNA fragments with diverse structures will compete for binding to Cas1-Cas2. To confirm the hypothesis that Cas1-Cas2 kinetically selects PS DNAs with long 3’-overhangs in a competitive environment, we designed an assay in which four different PS DNAs with distinct FRET efficiencies (Figure S1G) competed for binding to the surface immobilized Cas1-Cas2 molecules. When we incubated four PS DNAs with different 3’-overhang lengths (5-, 7-, 10-, and 15-nt-long) in equimolar ratios, the majority (95%) of the stably bound molecules had a non-canonical 3’-overhang, whereas the canonical PS DNA populated only 5% of the molecules (Figures 1J and S1G). Taken together, these results show that the Cas1-Cas2 complex effectively selects precursor PS DNAs with long 3’-overhangs in the cellular environment.

### Cas1-Cas2 Interacts Frequently with PAM-Containing Prespacers

Driven by the hypothesis that the frequent interactions of Cas1-Cas2 with long single-stranded 3’-overhangs may serve as a selection mechanism for suitable PS DNAs, we used the single-molecule assay to probe whether Cas1-Cas2 recognizes the PAM sequence during the short-lived interactions. First, we explored how the position of the PAM sequence in the 15-nt-long 3’-overhang affected the binding behavior of Cas1-Cas2. When the PAM sequence in 3’-overhang was moved away from the optimal position, fewer molecules were stably bound to the precursor PS DNA (Figures S2A and S2B), suggesting that Cas1-Cas2 probed the 3’-for PAM sequences. Consistent with this observation, recent structural studies showed that Gln287 and Ile291 in the proline-rich C-terminal tail of Cas1b recognize cytosine (C) at +5 and T at +6 of 5’-CTT-3’ PAM sequence in the 3’-overhang (Figure 1B, Figure S2C), resulting in stabilization of the C-terminal tail of Cas1b (Wang et al., 2015).

To further elucidate the PAM specificity of Cas1-Cas2, we generated 16 different precursor PS DNAs, which covered all the nucleotide combinations at positions of +5 and +6 of the optimally positioned PAM sequence (Figure 1B). In agreement with our hypothesis, Cas1-Cas2 interacted with the different PAM variants at distinct frequencies (Figures 2A and S2D), whereas the dissociation rate remained unaltered (Figures 2B). Among these substrates, Cas1-Cas2 showed the highest binding frequency for 5’-CTT-3’ PAM, suggesting that Cas1-Cas2 kinetically selects for PAM containing precursor DNAs. When we repeated the experiment with a C-terminal tail mutant (Cas1: Q287A/I291G) of Cas1-Cas2, the differences in the binding frequency among substrates was no longer observed (Figure 2C). This data suggests that the Cas1-Cas2 complex selects PAM containing precursor DNAs through frequent interactions that are mediated by specific amino acid residues in C-terminal tail of Cas1b.

**Figure 2:**
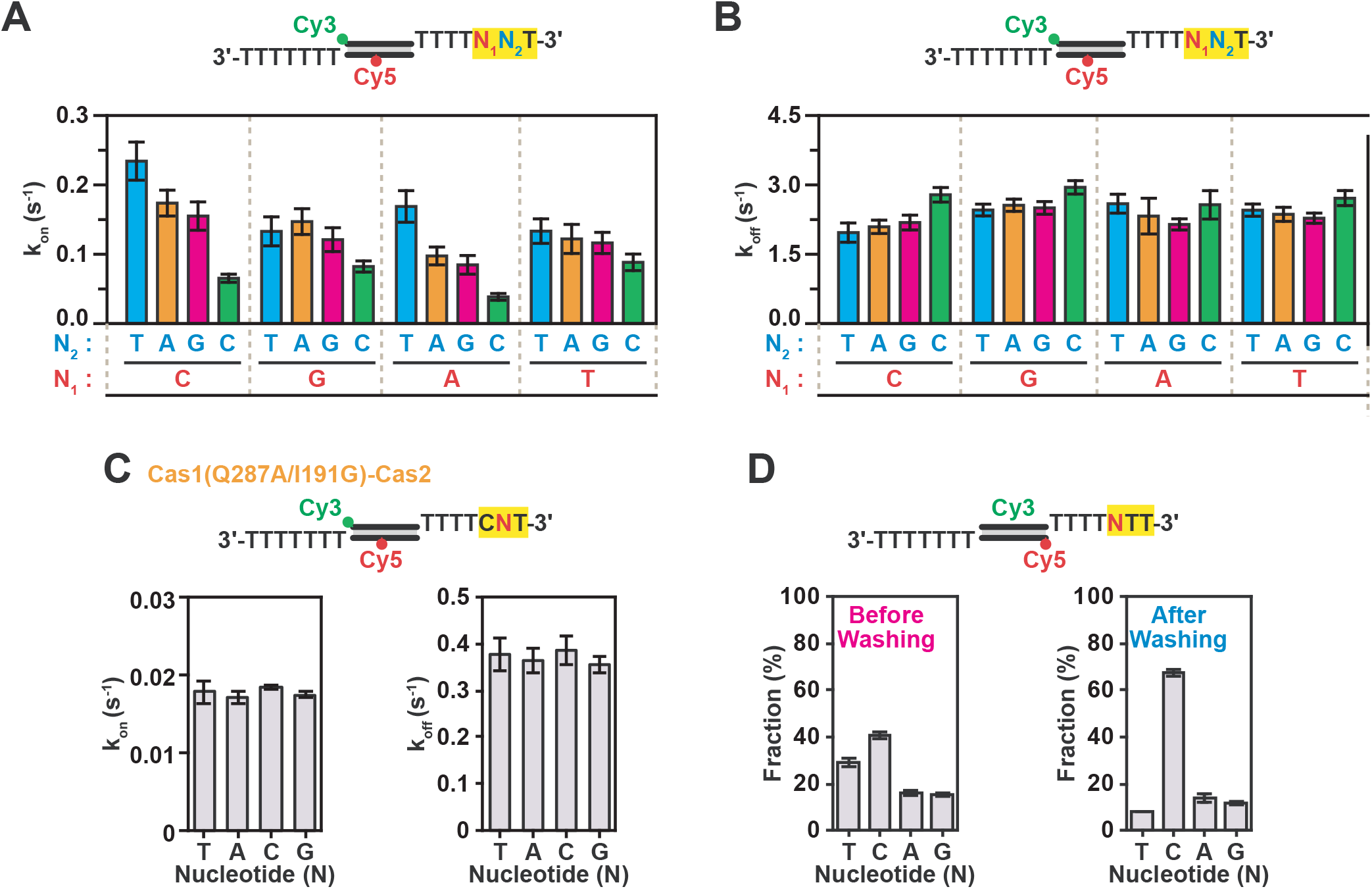
3’-Overhang Sequence-Dependent Prespacer Selection by Cas1-Cas2. **(A)** K_on_ values of short-lived binding events of various sequences at PS PAM recognition position with WT Cas1-Cas2. K_on_ was calculated in the same way as (Figure 1G). Error bars indicate the 95% confidence interval obtained by bootstrap analysis. **(B)** K_off_ values of short-lived binding events over various sequences at PS PAM recognition position with WT Cas1-Cas2. K_off_ was calculated in the same way as (Figure 1H). Error bars indicate the 95% confidence interval obtained by bootstrap analysis. **(C)** K_on_ and K_off_ values of short-lived binding for various sequences at PS PAM recognition position with mutant Cas1(Q287A/I291G)-Cas2. **(D)** Fractions of each FRET population of Cas1-Cas2-bound PS molecules various sequence PAM position sequences at 30 min after PS DNA addition and followed by washing to observe stably bound molecules. Data represents mean ± SEM (n = 3).

To investigate whether Cas1-Cas2 would preferentially bind precursor PS DNAs with 5’-CTT-3’ PAM in a competitive environment that mimics intracellular conditions, we employed our competition assay. When the Cas1-Cas2 complexes were presented with four different DNA substrates that contained different nucleotide identities at +5 position (Figure S2E), the 5’-CTT-3’ PAM was predominantly bound by Cas1-Cas2 compared to the other sequences at equilibrium (Figures 2D and S2F), thus, showing that Cas1-Cas2 complex kinetically selects for PAM-containing precursor DNAs. In conclusion, these experiments show that Cas1-Cas2 selects to PAM-containing PS precursors with a long 3’-overhangs through frequent interactions in competitive environments that mimic intracellular conditions.

### Cas1-Cas2 Facilitates Pairing of PAM-Containing Single-Stranded DNA

Recent studies have shown that the Cas1-Cas2 integration complex is able to repurpose DNA degradation fragments from RecBCD and Cas3 (Jackson et al., 2017; Kunne et al., 2016; Levy et al., 2015). Given the directionality and strand specificity of these enzymes, the degradation fragments are likely to be single-stranded when being released from the enzyme (Musharova et al., 2017). We hypothesized that Cas1-Cas2 might capture two complementary ssDNAs to form an effector integration complex. To visualize the potential interactions with ssDNA, we repeated the single-molecule assay with a series of ssDNA fragments (Figure 3A). When a ssDNA without PAM was introduced, no appreciable interactions were observed (Figure S3A). In contrast, when the ssDNA contained one or more PAMs, frequent transient interactions with the ssDNA were observed (Figure S3A). Interestingly, kinetic analysis of these binding events showed comparable binding frequency (Figures 3B and S3B) and dissociation rates (Figures 3C) among the substrates with a different PAM content. These results suggest that Cas1-Cas2 probes for PAM sites through random 3D collisions with ssDNA substrates.

**Figure 3:**
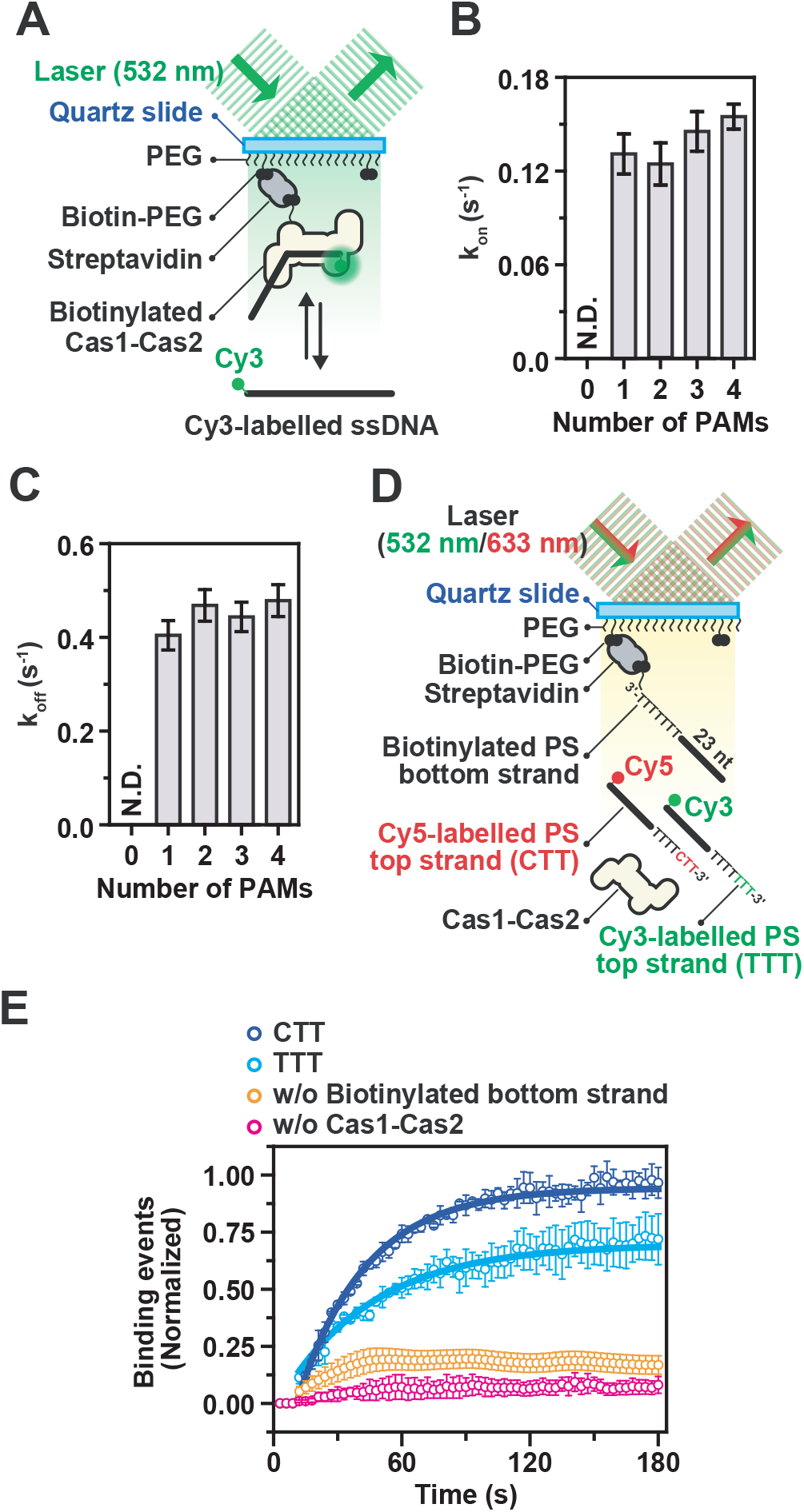
PAM-Dependent ssDNA Capture and Facilitated Loading by Cas1-Cas2. **(A)** Schematic representation of single-molecule assay for ssDNA binding of Cas1-Cas2. **(B)** The k_on_ values were calculated from cumulative probability of the arrival time for various numbers of PAM sequences in ssDNA. Error bars indicate the 95% confidence interval obtained by bootstrap analysis. **(C)** Dissociation rate (k_off_) of binding events of the various PAM numbers in ssDNAs. The koff values were calculated from dwell time (Δτ) of each event by single-exponential decay fitting. Error bars indicate the 95% confidence interval obtained by bootstrap analysis. **(D)** Schematic depicts of single-molecule assay for the detection of Cas1-Cas2-facilitated complementary strand loading. Cy3 was labelled on non-PAM (5’-TTT-3’) ssDNA strands and Cy5 on PAM (5’-CTT-3’) ssDNA strands. Biotinylated complementary strands were immobilized onto the surface of microfluidic slide chamber through biotin-streptavidin tethering. Green (532 nm) and red (633 nm) lasers were individually used to excite Cy3 and Cy5, respectively. **(E)** Normalized number annealing events to the surface immobilized complementary strand. Solid lines exhibit a single-exponential fit. Error bars represent the 95% confidence interval obtained through bootstrap analysis.

Given that PS DNAs need to be duplexed for spacer integration, we hypothesized that Cas1-Cas2 may facilitate recruitment of the complementary DNA strand, stabilizing the interaction. To investigate whether Cas1-Cas2 enhances the pairing between the PAM-containing top strand and its complementary bottom strand, we designed a single-molecule DNA capture assay (Figure 3D). In brief, the bottom strand without a PAM sequence was immobilized to the surface of a microfluidic chamber through biotin-streptavidin tethering. Next, two complementary top strands were introduced into the microfluidic chamber: a Cy5-labelled strand that contained the 5’-CTT-3’ PAM, and a Cy3-labelled strand that contained the 5’-TTT-3’ non-PAM in the optimal position of 3’-overhang (Figure 3D). In the presence of Cas1-Cas2, the number of top strands that bound to the immobilized bottom strands increased exponentially over time (Figure 3E). Moreover, the PAM-containing top strands accumulated at a faster rate than top strand with the non-PAM at the optimal position, resulting in more PAM-containing strands annealed. These results are consistent with electromobility shift assays (EMSA) (Figures S3C). Control experiments in the absence of Cas1-Cas2 or the biotinylated bottom strands showed no detectable accumulation of binding (Figure 3E). These results suggest that the Cas1-Cas2 complex facilitates the pairing ssDNA into precursor PS DNAs, in which the PAM sequence acts as essential identification marker.

### Identification of Prespacer 3’-Overhang Trimming Enzymes

Our single-molecule data shows that Cas1-Cas2 selects PAM-containing precursor PS DNA with long 3’-overhangs and promotes the formation of such precursor PS from ssDNA strands. However, the E. coli type I-E Cas1-Cas2 complex is only able to integrate canonical PS DNAs with 5-nt long 3’-overhang at both 3’-ends (Figure 4A) (Nunez et al., 2015b). Therefore, the PS precursors with long 3’-overhangs need to be matured into the canonical size of 5-nt for efficient integration into the CRISPR array. A recent structural study has suggested a cut-and-paste model in which the Cas1-Cas2 complex itself can mature a 3’-overhang into the optimal 5-nt size, using the endonucleolytic activity of Cas1 (Wang et al., 2015). However, using identical conditions, we could not reproduce this activity by the Cas1-Cas2 complex (Figures 4B and S4A), which suggests that there may be an alternative mechanism to produce canonical PS DNAs with 5-nt overhangs.

**Figure 4:**
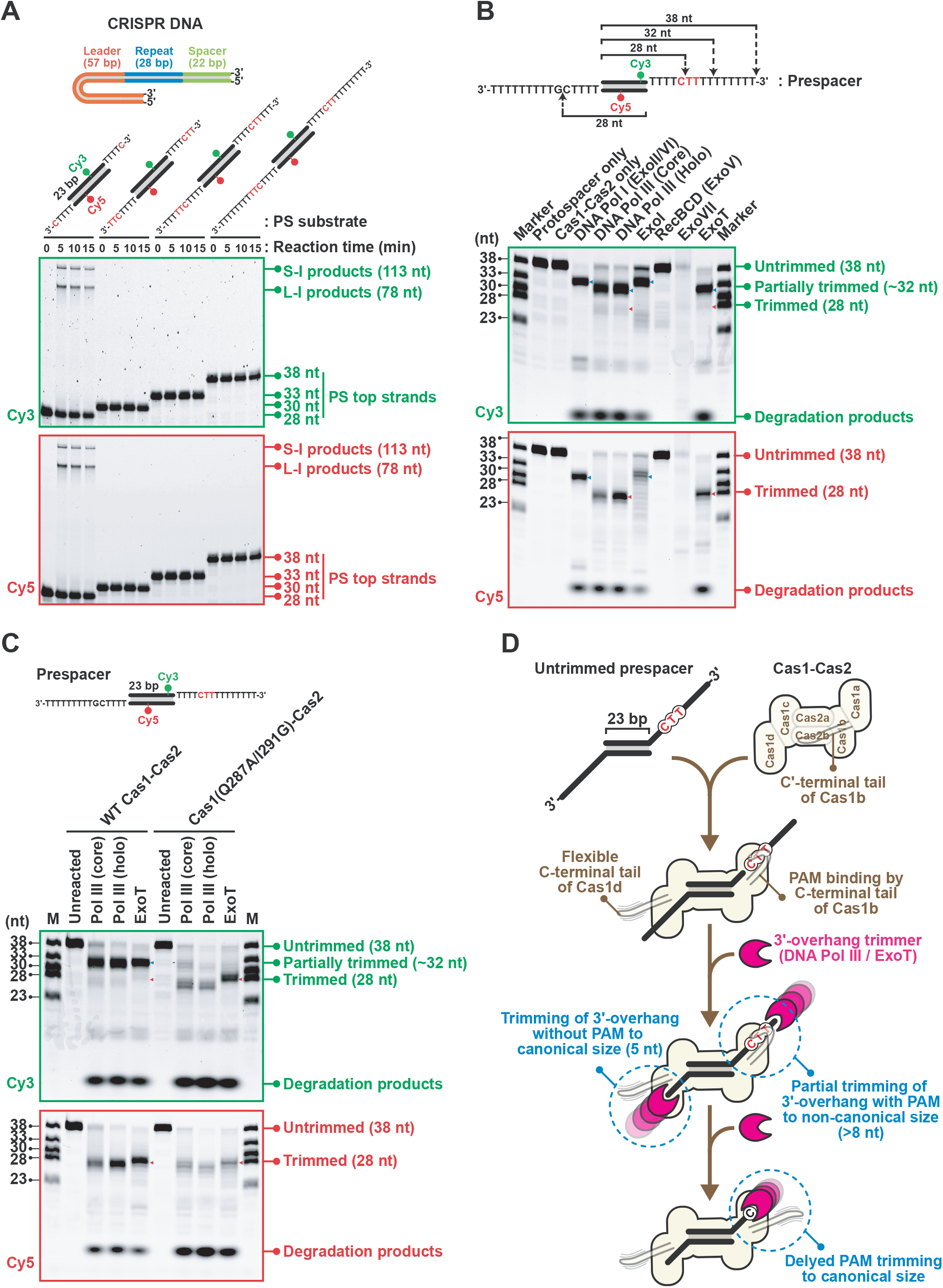
Identification of DNA Pol III and ExoT as PS 3’-Overhang Trimmers. **(A)** In vitro integration assay using PS DNAs with various 3’-overhang lengths. Full-site integration in CRISPR DNA (tandemly arrayed 57-nt leader, 28-nt-repeat and 22-nt partial spacer) produces 78-nt leader-side integration (L-I) products and 113-nt spacer-side integration (S-I) products in case of the use of PS DNA with 28-nt strands at both strands. Cy3 and Cy5 were labelled at the middle of both strands respectively as indicated. **(B)** In vitro trimming assay using wild type Cas1-Cas2 and various exonucleases with PS the represented DNA. Trimmed products (red) and partially trimmed products (blue) are indicated with triangles. Cy3- or Cy5-labelled 38-, 33-, 30-, 28- and 23-nt long ssDNAs were used and loaded as molecular size markers. **(C)** In vitro trimming assay using wild-type (WT) or mutant Cas1(Q287A/I291G)-Cas2 with/without core complex DNA polymerase III (Pol III (core)) or holoenzyme (Pol III (holo)) or Exonuclease T (ExoT). **(D)** Model for Cas1-Cas2 C-terminal tail-dependent protection of PAM in the 3’-overhang from trimming by DNA Pol III or ExoT.

A recent study on the CRISPR system in Streptococcus thermophilus reported that a Cas2 protein with a DnaQ-like domain exhibits trimming activity on PS precursor DNA, resulting in canonical PS DNA with 5-nt 3’-overhangs (Drabavicius et al., 2018). These fusions of Cas2 and DnaQ-like domains have been reported for several other type I-E systems (Horvath and Barrangou, 2010), hinting that proteins with DnaQ-like domains may naturally be essential for PS maturation. In search of PS trimming enzymes in the E. coli type I-E system, we identified replicative DNA polymerases as promising candidates. One of the crucial tasks of these polymerases (e.g. DNA Pol III) is proofreading for errors during synthesis, which requires 3’-5’ exonuclease activity (Hamdan et al., 2002; Huang et al., 1997). This 3’-5’ exonuclease activity in replicative DNA Pol III is achieved by its DnaQ ε subunit, which is a part of the core complex of the polymerase. Thus, we hypothesized that DnaQ of Pol III or other exonucleases with DnaQ-like domains might work as 3’-overhang trimming enzymes (here after called trimmers) to generate functional PS DNA in the E. coli type I-E CRISPR-Cas system.

To test the hypothesis if nucleases with the 3’-5’ exonuclease activity or DnaQ-like domains are capable of processing precursor PS DNAs, we tested the following enzymes: DNA Pol I (ExoII/VI), DNA Pol III core complex (αεѳ subunits), DNA Pol III holoenzyme (holo), Exonuclease I (ExoI), RecBCD (ExoV), Exonuclease VII (ExoVII) and ExoT (Lovett, 2011). To observe the trimming activity of these enzymes, we designed an in vitro trimming assay in which the Cy3-labelled top strand of the PS precursor contained a PAM (5’-CTT-3’) and the complementary Cy5-labelled bottom strand of the PS precursor contained non-PAM (5’-CGT-3’) (Figures 4B). This scheme enabled us to distinguish the efficiencies of 3’-5’ exonuclease enzymes on both PAM- and non-PAM-containing strands of the Cas1-Cas2 bound PS precursor.

The in vitro trimming assay revealed that DNA Pol I and ExoI exhibited a weak level of trimming, as a result the majority of products were partially trimmed (~32 nt) and cannot be used as substrate for integration by Cas1-Cas2 (Figure 4B). Interestingly, DNA Pol III core complex, holoenzyme and ExoT could trim the 3’-overhang of the PAM-deficient bottom strand to the canonical size 28-nt (5-nt 3’-overhang, Figures 4B and S4A). In contrast to the bottom strand that lacked a PAM sequence, the PAM-containing top-strand was partially trimmed to a non-canonical size of (~31 nt) (Figures 4B and S4A). When the assay was repeated with a PS precursor without any PAM sequence, both strands were trimmed equally into the canonical size 28-nt (Figure S4B). In contrast, a PS precursor with a PAM sequence in both strands showed trimming to a non-canonical size of (~31 nt) (Figure S4B). These results demonstrate that PS precursors with only one PAM in the 3’-overhangs are asymmetrically trimmed by DNA Pol III and ExoT.

### Cas1-Cas2 Protects PAM in the Prespacer 3’-Overhang from Trimming

Our single-molecule assay revealed that the C-terminal tail of Cas1 is essential for distinguishing the nucleotides of the PAM sequence (Figure 2C), which may hint towards mechanism for the asymmetric trimming of PS precursors. Consistent with this hypothesis, the crystal structure of Cas1-Cas2 in complex with a PAM-containing 7-nt-long 3’-overhang-containing PS DNA shows that the C-terminal tail of Cas1b protects the PAM site in 3’-overhangs from nuclease attacks via stable interactions with the PAM (Figure S2C) (Wang et al., 2015). In contrast, the electron density for the C-terminal tail of Cas1b was absent in the crystal structure of Cas1-Cas2 in complex with a PAM-deficient precursor prespacer (Figure S2C) (Wang et al., 2015). Therefore, we hypothesized that the Cas1b C-terminal tail-mediated PAM recognition might protect the PAM-sequence from trimming, resulting in asymmetrically trimmed PS DNAs.

To test this hypothesis, we repeated the in vitro trimming assay with WT or C-terminal tail mutant of Cas1 (Q287A/ I291G) in the Cas1-Cas2 complex. Consistent with our previous results, both DNA Pol III core complex and ExoT trimmed the precursor PS bound to WT Cas1-Cas2 asymmetrically. However, in the case of C-terminal tail mutant, DNA Pol III core complex and ExoT trimmed both strands into the canonical size of 28-nt, regardless of the presence of a PAM sequence (Figures 4C and S4C). From these experiments, we conclude that the C-terminal tail of Cas1b in Cas1-Cas2 protects the PAM sequence during maturation of PS precursors, resulting in PS DNA with asymmetrically trimmed 3’-overhangs (Figure 4D).

### Asymmetric Trimming of PS Precursors Results in Biased Leader-Side Integration

Previous studies have shown that in order to obtain functional spacers, the PAM-deficient PS strand must be integrated at the leader-side (L-site) of CRISPR-array, commonly referred to as the half-site intermediate (Nunez et al., 2016; Wright et al., 2017). Subsequently, the PAM-containing strand is integrated at spacer-side (S-site) of the CRISPR-array to complete the integration process (Nunez et al., 2016; Wright et al., 2017). We hypothesized that the asymmetric trimming of precursor PS may serve as a mechanism to ensure that the canonical PAM-deficient strand (28 nt) of the PS is integrated first at the L-site, followed by integration of the non-canonical PAM-containing strand (31 nt) at the S-site if the PAM-containing strand is trimmed after L-site integration. Such a mechanism would heavily bias integration for correctly oriented spacers.

To test if the asymmetric trimming of a precursor PS would generate a bias for correctly oriented spacers, we repeated the in vitro integration assay and sought to verify whether the first integration event only occurs at the L-site when the Cas1-Cas2 was bound to an asymmetrically trimmed PS DNA. When the Cas1-Cas2 complex was bound to various mimics of asymmetrically trimmed PS DNA, the integration of the trimmed PAM-deficient strands was only observed at the L-site (Figure 5A), supporting the model of stepwise integration in the CRISPR-array (Xiao et al., 2017). In contrast, control experiments with a canonical PS that has already been matured resulted in both S-site and L-site integration (Figure 5A), suggesting that the bias disappeared.

**Figure 5:**
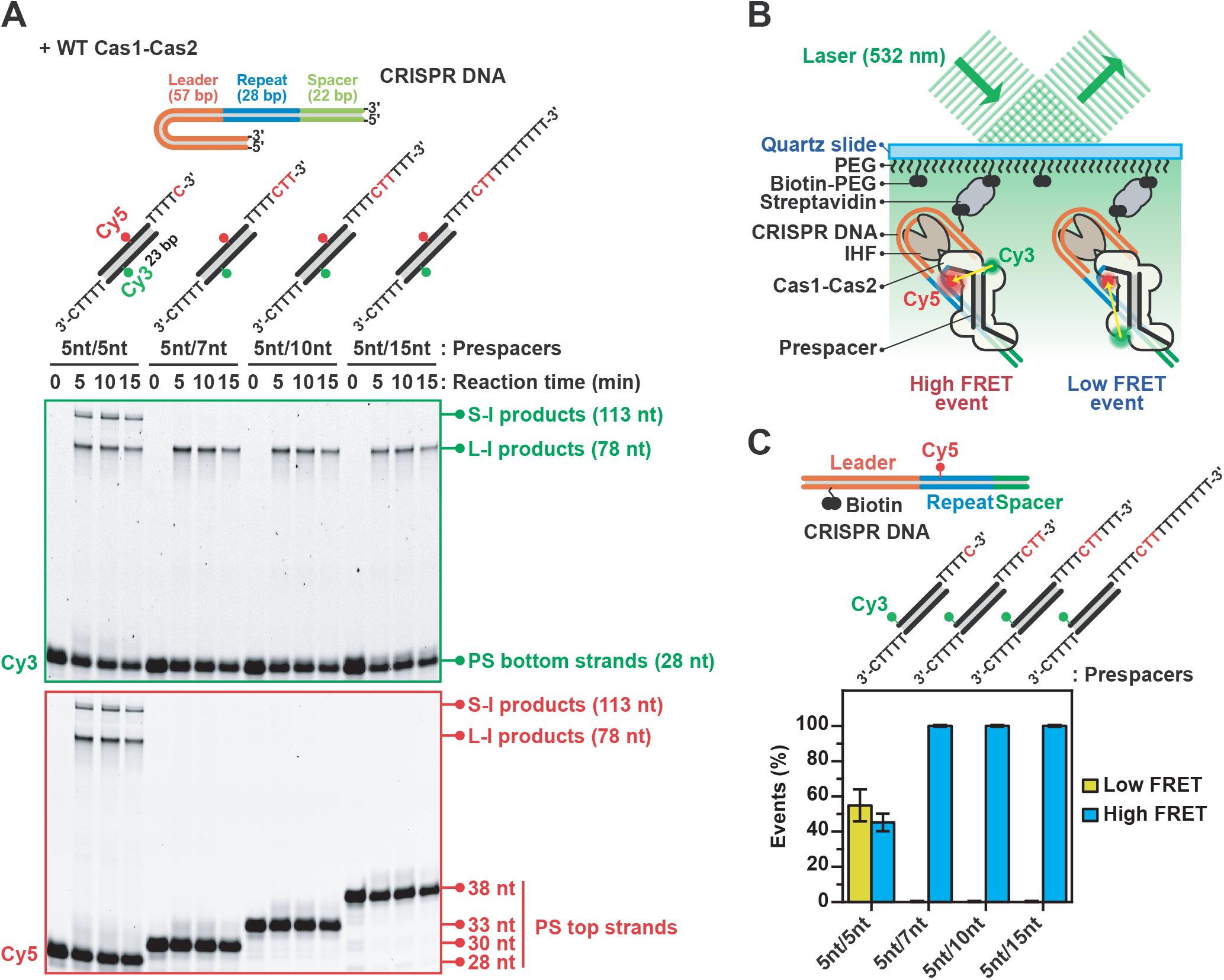
Leader-Proximal Integration of Non-PAM-Derived Ends of Asymmetrically Trimmed PS DNA. **(A)** In vitro integration assay using PS DNAs with various 3’-overhang lengths in top strands but with optimally trimmed length (28-nt) in bottom strands. The assay was performed as same in Figure 3A. **(B)** smFRET design to probe the orientation of integrated products. 5’-end of top strand was labelled with Cy3. Integration of 3’-end of bottom strand in leader-side integration site exhibits high FRET, and integration of 3’-end of top strand shows low FRET. **(C)** Event percentages of high and low FRET populations. Design of biotinylated CRISPR DNA, which was labelled with Cy5 in repeat at 5-nt away from leader-repeat junction, is presented. PS DNAs used in smFRET assay are presented above bar graphs. Data is represented as mean ± SEM (n = 3).

To validate the biochemical assays, we designed an assay based on smFRET to track spacer integration at the single-molecule level (Figure S5A). In brief, a Cy5-labelled CRISPR-array (Cy5) was immobilized to the surface of a microfluidic chamber. Next, the integration host factor (IHF) protein, which is critical for the chronological integration of new spacers at the leader side of the CRISPR-array (Figure S5B) (Nunez et al., 2016; Wright et al., 2017; Yoganand et al., 2017), was incubated with the surface-immobilized CRISPR-array. After washing, Cas1-Cas2 bound to a labelled PS DNA (Cy3) was added to the slide, and integration events were tracked by smFRET. In this scheme, L-site integration of the PAM-deficient strand will result in high FRET, whereas S-site integration events of the PAM-deficient strand will result in low FRET (Figures 5B and S5C). Consistent with our previous biochemical assays, canonical PS DNA resulted in a 50/50 ratio for L-site and S-site integration (Figures 5C and S5D). In contrast, the asymmetrically trimmed PS precursors exclusively populated high FRET, suggesting that these substrates were integrated only in the L-site of the CRISPR array (Figures 5C and S5D). Taken together, these results suggest that asymmetric trimming of PS precursors creates a bias for spacer integration in the correct orientation.

### Delayed PAM Trimming Leads to Correctly Oriented Spacer Integration

The data above show that asymmetric trimming of PS precursors generates a bias for integration of the PAM-deficient strand of the PS at the L-site of the CRISPR-array. To complete integration, the PAM-containing end of the PS precursor must also be trimmed to a canonical size of 28 nt (5-nt overhang) for integration at the S-site of the CRISPR array. To determine whether DNA Pol III (core) and ExoT can process the PAM-containing partially trimmed 3’-overhangs, we performed in vitro trimming-driven integration assays with half-site intermediate DNA constructs (Figure S6A). These substrates comprised PAM-containing upper PS strands with a 7, 10, and 15-nt long non-canonical 3’-overhang (Figure 6A and S6B). Initial experiments showed that the 3’ ends of the oligonucleotide based CRISPR-array were non-specifically degraded by the 3’-5’ exonuclease activity of DNA Pol III (core) and ExoT (Data not shown). To prevent this non-specific degradation at both 3’-ends of CRISPR array DNA were modified with three consecutive phosphorothioate (PTO) backbone linkages (Figure S6A).

When DNA Pol III or ExoT was incubated with Cas1-Cas2 in complex with the half-site intermediate constructs, the trimming enzymes were able to mature the PAM-containing top strand of PS DNA to a canonical size, resulting in distinct S-site integration products (Figures 6A and S6B). Interestingly, the overall efficiency of the full-site integration declined with an increase of the length of PAM-containing overhang. This suggests that substrates that are close to the optimal size are integrated at higher efficiencies compared to substrate that require more extensive trimming (Figures 6A and S6B). In conclusion, these experiments show that each of DNA Pol III and ExoT drives full-site integration of unprocessed half-site intermediate PS DNAs with non-canonical 3’-overhangs.

**Figure 6:**
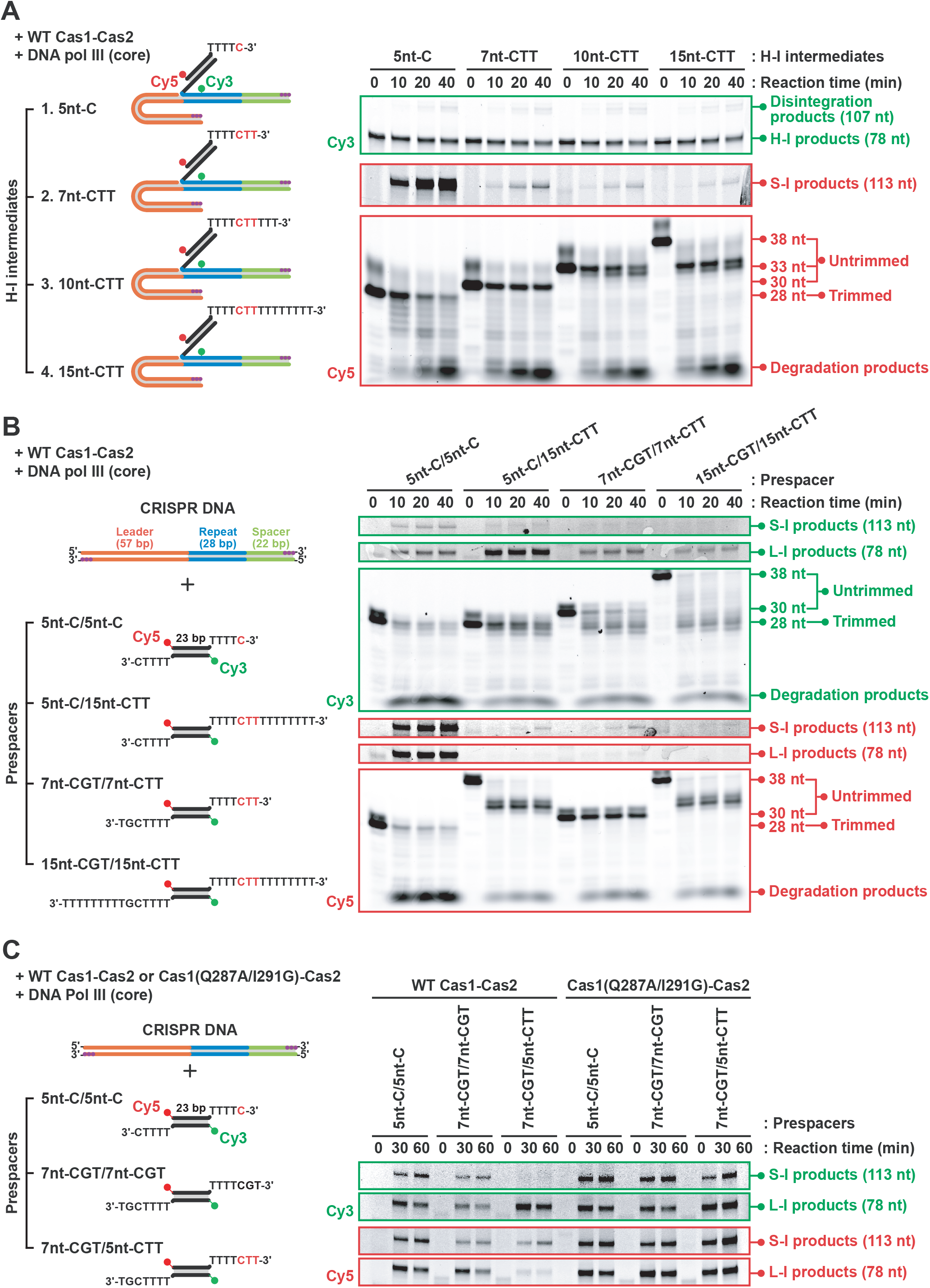
Delayed PAM Trimming Results in Correctly Oriented Spacer Integration. **(A)** Gel images of in vitro trimming-driven full-site integration assay with DNA Pol III (core). Unreacted half-site intermediates, disintegrated products are presented in the Cy3 image. Spacer-side integration products and processed top PS strands are represented in the Cy5 images. To clearly exhibit, the bottom part (below 50 nt) of the Cy5 image was separated from the top part (around 70~130 nt) and adjusted with different contrasts. **(B)** Representative gel images of in vitro trimming-driven integration assay with DNA Pol III (core). Contrast of areas of spacer-side and leader-side integration products were adjusted for the optimal visibility. **(C)** Wild type (WT) or C-terminal end mutant (Q287A/I291G) Cas1-Cas2 were used for in vitro trimming-driven integration assay with DNA Pol III (core). Dye-labelled canonical PS DNA (5nt-C/5nt-C) or non-canonical PS DNA without PAM (7nt-CGT/7nt-CGT) or with PAM (7nt-CGT/7nt-CTT) were used in the assay. Linear CRISPR DNA substrate was modified by three consecutive PTO at the both ends to be protected from nonspecific degradation by DNA Pol III. Leader-side integration (L-I) and spacer-side integration (S-I) products exhibit as 78 nt and 113 nt sizes, respectively.

Next, we investigated whether the Cas1-Cas2 and IHF dimer together with the PS trimmer, DNA Pol III (core), are sufficient to obtain correctly orientated spacer integration of PS precursors in vitro. To observe trimming-driven integration, we performed in vitro trimming-driven integration assays with a PTO-modified linear CRISPR-array and various combinations of PS DNAs. These PS DNAs included: a canonical PS (5-nt), a partially trimmed substrate (7-nt) and an untrimmed (15-nt) substrate, which contained either a PAM (5’-CTT-3’) or non-PAM (5’-CGT-3’) sequence at each strand (Figures 6B and S6C). Consistent with our previous data, the canonical PS DNA with a 5-nt 3’-end (5nt-C/5nt-C) resulted in a comparable ratio of correctly and incorrectly oriented full-site integration products. However, when the assay was repeated with an asymmetrically trimmed (5nt-C/15nt-CTT), partially trimmed (7nt-CGT/7nt-CTT) and untrimmed (15nt-CGT/15nt-CTT) PS DNA, the full-site integration products were biased towards the S-site of the CRISPR-array. These results confirm our previous trimming data where PAM-deficient 3’-overhangs are trimmed to an optimal size for integration (28-nt) and PAM-containing 3’-overhangs are partially trimmed (~31-nt, Figures 6B). Notably, the canonical PS DNA with 5-nt 3’-overhangs was more excessively degraded by DNA Pol III (core), suggesting that fewer Cas1-Cas2 molecules were stably bound to the PS DNA (Figure 6B). This agrees with our previous single-molecule binding data (Figures 1C and 1I), which showed that the number of stably bound molecules increased proportional with the 3’-overhang length.

To confirm the trimming-driven integration data, we further explored the integration behavior using smFRET. To probe integration of PS DNAs at the single-molecule level, we incubated the biotinylated, Cy5 labelled and PTO-modified CRISPR array DNA constructs with Cas1-Cas2 proteins that were pre-incubated with PS substrates. And then we treated with DNA Pol III (core), prior to immobilization onto the surface of a microfluidic chamber (Figure S6D). Next, we collected the FRET efficiencies of single-molecules (Figure S6E). Consistent with our biochemical observations, the integration of trimmed bottom 3’-end of a PS DNA with a PAM sequence in the upper strand was biased to the L-site of the CRISPR-array, resulting in high FRET (15-nt-TTT/15nt-CTT). In contrast, the trimmed 3’-ends of a PS DNA with two non-PAM containing 3’-overhangs (15-nt-TTT/15nt-CGT) resulted in both L-site (high FRET) and S-site (low FRET) integration, albeit at different efficiencies (Figure S6E). Notably, these results suggest that PAM-like sequences with a C at +5 of the PS DNA are more efficiently stabilized than other sequences, which is in good agreement with our single-molecule binding and EMSA data (Figure 2A and S1F).

To clarify the role of PAM protection by Cas1 C-terminal tail in this orientation bias, we repeated in vitro trimming-driven integration assay with the C-terminal mutant of Cas1-Cas2 Cas1(Q287A/I291G)-Cas2. In contrast to wild type Cas1-Cas2 that resulted in a strong bias for correctly oriented integration of PAM-containing PS, the C-terminal mutant Cas1-Cas2 complex exhibited unbiased integration of PAM-containing PS DNA (7nt-CGT/7nt-CTT), similar to canonical PS (5nt-C/5nt-C) and non-canonical but PAM-deficient PS (7nt-CGT/7nt-CGT) substrates (Figure 6C). We speculate that this spectrum in PAM stabilization facilitates a robust spacer integration reaction. In conclusion, we show that delayed PAM trimming of precursor PS DNAs results in a strong bias for correctly oriented spacer integration in the CRISPR array, conferring robustness to CRISPR-Cas immunity (Figure 7).

**Figure 7:**
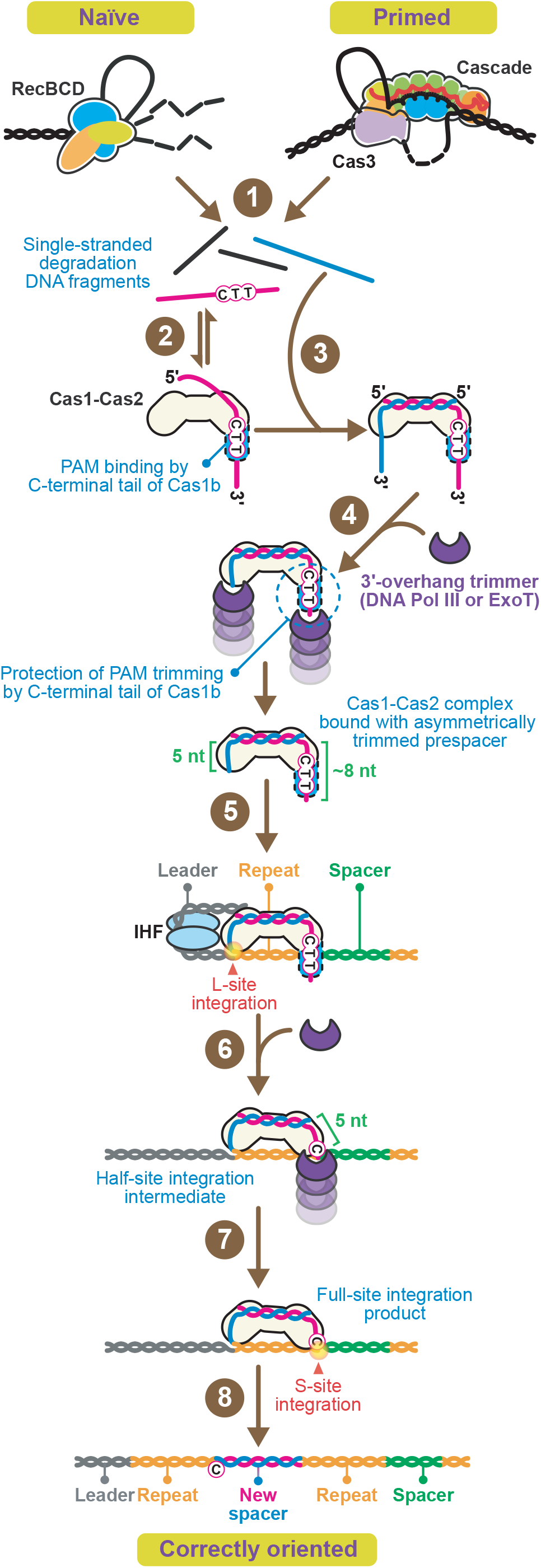
Model for PS Selection, Trimming and Correct Orientation of Integrated Spacers. **(1)** RecBCD and Cascade-Cas3 complexes generate single-stranded degradation DNA fragments during naïve and primed adaptation. **(2)** The PAM-containing (5’-CTT-3’) ssDNA are captured and annealed by Cas1-Cas2. **(3)** Complementary strand is base-paired. Consequently, most of the prespacer DNAs loaded onto Cas1-Cas2 have a PAM at +5 to +7 in one of 3’-overhang that is be longer than 7 nt. **(4)** DNA Pol III or ExoT asymmetrically trim 3’-overhangs of the PAM-deficient strand to the canonical size (5 nt), whereas the PAM-containing strand is partially matured (~7 nt). **(5)** The non-PAM-derived 3’-end is subject to the first integration event that only occurs at the leader-side of the first repeat. **(6)** The partially trimmed PAM-derived 3’-overhang of the half-site intermediate is further trimmed into canonical size (5 nt). **(7)** Subsequently, this PAM-derived end is integrated in spacer-side integration site. **(8)** Finally, host DNA repair enzymes fill the integration site and repeat duplication occurs eventually. In contrast to conventional unbiased trimming model (Figure S7), this “delayed PAM trimming” model results in strong bias for correct orientation of newly integrated spacer in the CRISPR array.

## Discussion

Prokaryotes harbor CRISPR-Cas adaptive immune systems to eliminate invading phages and mobile genetic elements. A key step towards immunity is the formation of immunological memory by the Cas1-Cas2 integration complex, which allows precise programming of effector complexes to their invading nucleic acid targets. Despite several seminal studies using biochemistry and cell biology approaches, many of the molecular details have remained elusive due to the lack of a method to track spacer integration in real-time. Here, we use a combination of single-molecule fluorescence spectroscopy and biochemistry, to understand the molecular details of the capture, processing and integration of DNA substrates by the E. coli Cas1-Cas2 integration complex.

Here we propose a model in which Cas1-Cas2 captures a wide variety of DNA substrates, including ssDNA and partially duplexed precursor PS with long 3’-overhangs (Figure 7). Guided by the PAM sequence, Cas1-Cas2 kinetically select prespacers that are suitable for integration in the CRISPR array. Once loaded, the long 3’-overhang of the precursor PS DNA is trimmed by DNA pol III and other DnaQ-like proteins. While the PS becomes matured by these enzymes, the PAM is protected by the C-terminal tail of Cas1, resulting in asymmetric maturation of the PS DNA. This asymmetry in maturation, generates a bias for leader side integration of the non-PAM end of the PS. Once leader side integration has occurred, the PAM end of the PS is released from Cas1, subsequently matured and integrated, resulting in correctly oriented and functional spacers in the CRISPR array.

### Cas1-Cas2 May Selectively Obtain Precursor Prespacers from Cellular DNA Sources

Our kinetic study shows that Cas1-Cas2 dynamically interacts with PS precursors DNA via frequent short-lived interactions. The frequency of the interactions is determined by sequence and length of the 3’-overhang of the PS precursor. The probability of transitioning from weakly associated status to the stable Cas1-Cas2-PS DNA complex is higher for PS DNA that has a longer ssDNA 3’-overhang and a 5’-CTT-3’ PAM at position 5 nt away from the end of duplex. These data are in good agreement with a recent molecular dynamics simulations study, which suggest that the interaction between Cas1-Cas2 and the PS DNA is stabilized when a PAM is present in the 3’-overhang (Wan et al., 2019). Moreover, our data shows that sequence specificity of Cas1-Cas2 is lost when the C-terminal tail of Cas1 is mutated, suggesting that Cas1 initially probes the 3’-overhangs of the precursor for PAM sequences, before it transitions into a stably bound state. Moreover, we show that in competitive environments, Cas1-Cas2 selects PAM-containing precursor PS that can subsequently be matured into functional spacers. This is consistent with in vivo data that show newly acquired spacers are preferentially obtained from PAM flanking sites (Savitskaya et al., 2013).

Our single-molecule assay with ssDNA substrates revealed that Cas1-Cas2 can capture PAM-containing ssDNA fragments and facilitate the formation of a precursor PS by recruiting the complementary strand. Based on these finding, we propose a new model, in which Cas1-Cas2 captures ssDNA fragments in the cell to acquire functional PS substrates. In this model, Cas1-Cas2 transiently interacts with the ssDNA through random 3D collisions to search for a PAM sequence. Next, it forms a stable effector complex by recruiting a complementary DNA. Such a mechanism would allow Cas1-Cas2 to repurpose the ssDNA degradation products of RecBCD and Cascade-Cas3. Interestingly, Cas1-Cas2 has been shown to form a complex with Cascade-Cas3 during primed spacer acquisition (Dillard et al., 2018; Redding et al., 2015). We hypothesize that this complex formation allows direct transfer of the ssDNA fragments to Cas1-Cas2, ensuring robust primed spacer acquisition. Recent single-molecule studies have shown that Cas3 generates ssDNA loops during CRISPR interference (Dillard et al., 2018; Loeff et al., 2018). It will be of interest to find out whether this ssDNA loop marked with PAM sequences can be directly recognized by the Cas1-Cas2 complex.

### DnaQ Exonucleases Mature Prespacers in Concert with Cas1-Cas2

The binding of precursor PS requires maturation of the 3’-overhang in order to be integrated into the CRISPR array. Recent structural studies speculated that Cas1-Cas2 itself could process the 3’-overhangs of precursor PS DNA, by using the residues that are involved in nucleophilic attack of the scissile 3’-ends during integration process (Wang et al., 2015). However, here we show that Cas1-Cas2 does not have 3’-overhang trimming activity in vitro (Figures 4B and S4A). Instead, we show that the 3’-5’ exonuclease activity of DNA pol III and ExoT allow for the maturation of precursor PS DNA.

DNA Pol III is the primary enzyme complex that is involved in DNA replication and is highly conserved among prokaryotes. The ε subunit, or DnaQ, lays at the core of this complex and facilitates proofreading during replication by means of its 3’ to 5’ exonuclease activity. Given that the majority of newly acquired spacers arise from stalled replication forks (Levy et al., 2015), we speculate that the stalled DNA Pol III might directly mature RecBCD degradation products bound by Cas1-Cas2. Moreover, the fact that DNA Pol III has DNA polymerization activity hints towards a molecular link between PS trimming and repeat duplication through trimming-driven PS integration.

The intracellular copy numbers of DNA Pol III holoenzyme are low (10–20 copies per cell) and its expression level is tightly regulated (Wu et al., 1984). This necessitates the involvement of abundant 3’-5’ exonucleases for efficient precursor PS processing, allowing the CRISPR-Cas immune system to keep pace with rapid infections. ExoT is an abundant and robust 3’-5’ exonuclease, ensuring high fidelity in DNA repair pathways (Hsiao et al., 2014; Viswanathan et al., 1998). The contribution of ExoT may compensate for the limited availability of DNA Pol III, during PS maturation. In addition, it is likely that several other unidentified exonucleases may also be involved in PS maturation.

### Delayed PAM Trimming Determines the Orientation of New Spacers

To date, the general working model of the spacer acquisition described PS binding, maturation and integration as independent steps (Figure S7). This model was incomplete, as it necessitates symmetrically trimmed canonical PS DNA for integration, which cannot show how the 3’-end cytosine of the PAM is correctly oriented for spacer integration. Here we show that the process of PS binding, maturation and integration is tightly coordinated in time. In our model, the PAM sequence of the prespacer is initially protected by the C-terminal tail of Cas1 from maturation, resulting in an asymmetry in PS trimming. The non-PAM end of the asymmetrically trimmed PS is first integrated at the leader side of the CRISPR-array, resulting in a half side intermediate. This integration step is followed by the release, maturation and integration of the PAM-end of the PS at the spacer side of the CRISPR-array. This delayed PAM trimming model explains how newly acquired spacers can be correctly oriented to generate functional spacers with high-fidelity. Given that Cas1-Cas2 is highly conserved, we anticipate that this mechanism of delayed PAM trimming can be generally applied to other CRISPR-Cas systems.

### Outlook

Recently, several groups have harnessed the nucleic acid acquisition capabilities of Cas1-Cas2 to develop new types of nucleic acid recording techniques in cellular contexts (Schmidt et al., 2018; Sheth and Wang, 2018; Sheth et al., 2017; Shipman et al., 2016, 2017). Cas1-Cas2-based recording techniques allow for capturing cellular events in prokaryotes in a chronical order (Schmidt et al., 2018; Sheth and Wang, 2018; Sheth et al., 2017; Shipman et al., 2016, 2017). Our findings may contribute to develop next generation of Cas1-Cas2 recorders with the increase in the density and accuracy of information storage. We anticipate that these findings may help to develop Cas1-Cas2-based recording system in eukaryotes, which has not been reported to date.

## Supporting information

Supplementary Information Kim & Loeff et al. April 2019

## Supplemental information

Supplemental information includes six figures (Figure S1–7) and one table (Table S1).

## Acknowledgements

We thank Anna Haagsma and Tim Künne for providing Cas1-Cas2 vectors and protein purification, the members of Joo and Brouns laboratories for discussions. We appreciate Samuel Leachman and Nynke Dekker (Delft University of Technology, the Netherlands), and Nicholas Dixon (University of Wollongong in Australia) for providing DNA polymerase III proteins. C.J. and S. J. J. Brouns were funded by the Foundation for Fundamental Research on Matter (Projectruimte 15PR3188). S. K. was partly funded by the European Union’s Horizon 2020 research and innovation programme under the Marie Skłodowska-Curie Grant Agreement No. 753528.

## Author Contributions

S.K., L.L., S.J.J.B. & C.J. conceived the study. S.K. and S.C. purified and labelled Cas1-Cas2 proteins and performed biochemical experiments. S.K. and L.L. performed single-molecule TIRF microscopy and analyzed data. S.K, L.L., S.J.J.B. and C.J. discussed the data and wrote the manuscript.

## Methods

### Protein Preparation

Cas1-Cas2 complex was expressed in E. coli BL21-AI chemically competent cells (Invitrogen™, Cat# C607003) using plasmids listed in Table S1 and purified using the N-terminal StrepII tag on Cas1 as described (Kunne et al., 2016). Briefly, cells were grown to an OD_600_ of 0.4 in LB media, cooled on ice for 30 min and induced with 0.5 mM IPTG and 0.2% L-arabinose. Protein expression was induced overnight at 20 °C. Cells were collected by centrifugation and lysed in 20 mM HEPES-NaOH pH 7.5, 75 mM NaCl, 1 mM DTT, 5% glycerol, 0.1% Triton X-100 using a Stansted pressure cell homogenizer. The lysate was cleared by centrifugation and incubated with Strep-Tactin beads (IBA) for 1 hour at 4 °C. Next, the lysate with beads were loaded onto a gravity column and washed with 20 mM HEPES-NaOH pH 7.5, 300 mM NaCl, 1 mM DTT, 5% glycerol, followed by elution with a buffer that contained 20 mM HEPES-NaOH pH 7.5, 75 mM NaCl, 1 mM DTT, 5% glycerol (Storage buffer) with 4 mM D-desthiobiotin (Cat# 2-1000-005). The presence and purity of the Cas1-Cas2 complex was checked through Bis-Tris 4-12% NuPAGE (Thermo Scientific, Cat# NP032A) with NuPAGE™ MES SDS Running Buffer (Thermo Scientific, Cat# NP0002). Protein complex was concentrated with Amicon^®^ Ultra Centrifugal Filters (Merck Millipore) and further purified on a Superdex 200 10/300 GL size-exclusion column (GE Healthcare, Cat# 17517501) using the ÄKTA pure protein purification system (GE Healthcare). The final complex was diluted in Storage buffer with 50% glycerol, snap-frozen in liquid nitrogen and stored at −80 °C. Mutant Cas1-Cas2 complexes were cloned by site-directed mutagenesis as described previously (Kim et al., 2015) and purified as described above. Primers for sub-cloning are listed in Table S1. The ihf-α and ihf-β genes were PCR amplified from E. coli (BL21) genomic DNA using the indicated primers in Table S1 and subsequently cloned into Berkeley MacroLab ligation-independent cloning (LIC) vectors using either LIC into 13K-HR and 13S-A vectors respectively, as described previously (Kieper et al., 2018). The ihf-α gene was cloned as an N-terminal His6 tag. Subsequent steps for the purification of IHFα/β dimer complex was performed as previously described (Nunez et al., 2016).

For site-specific Cas1-Cas2 biotinylation, an N-terminal LCTPSR FGE recognition motif on Cas1 was inserted by site-directed mutagenesis in plasmid pWUR871 with primers in Table S1 and co-expressed with FGE proteins (Addgene, plasmid #16132) (Carrico et al., 2007). Purified Cas1-Cas2 complex was buffer-changed with 0.5 M Sodium acetate (pH 5.5) and labelled with EZ-Link™ Biotin-LC-Hydrazide (Thermo Scientific, Cat# 21340) and incubated overnight at room temperature. Labelled Cas1-Cas2 was purified by size exclusion chromatography with Superdex 200 10/300 GL. Fractions were concentrated using Amicon^®^ Ultra Centrifugal Filters, pooled in Storage buffer with 50% glycerol, snap-frozen in liquid nitrogen and stored at −80 °C.

### DNA Preparation

Synthetic DNA oligonucleotides (Ella Biotech) were internally labelled with a monoreactive NHS-ester form of cyanine dyes as donors (Cy3, GE Healthcare, Cat# PA13101) or acceptors (Cy5, GE Healthcare, Cat#PA15101) or EZ-Link™ NHS-Biotin (Thermo Scientific, Cat# 20217) at amino-C6-dT (amine modification with amino-modifier C6-T) (Table S1). After labelling, the ssDNA strands were annealed in 20 mM Tris (pH8.0), 150 mM KCl and 5 mM MgCl_2_ using a thermocycler (Bio-Rad) at −1 °C/cycle for 1 min/cycle from 95 °C to 16 °C and then store at 4 °C.

### Single-Molecule TIRF Imaging and Data Acquisition

The fluorescent label Cy3 and Cy5 were imaged using prism-type total internal reflection microscopy (Loeff et al., 2018). In brief, Cy3 was imaged through excitation by a 532 nm diode laser (Compass 215M-50, Coherent). Cy5 was detected via FRET with Cy3, but if necessary, Cy5 was directly excited using a 640 nm solid-state laser (CUBE 640-100C, Coherent). Fluorescence signals from single molecules were collected through a 60x water immersion objective (UPlanSApo, Olympus) with an inverted microscope (IX71, Olympus). Scattering of the 532 nm laser beam was blocked with a 550 nm long-pass filter (LP03-532RU-25, SemRock). When the 640 nm laser was used, 640nm laser scattering was blocked with a notch filter (633 ± 12.5nm, NF03-633E-25, SemRock). Subsequently, signals of Cy3 and Cy5 were spectrally split with a dichroic mirror (λ_cutoff_ = 645 nm, Chroma) and imaged onto halves of an electron multiplying CCD camera (iXon 897, Andor Technology).

To eliminate non-specific surface adsorption of proteins and nucleic acids to a quartz surface (Finkenbeiner), pira-nha-etched slides were PEG-passivated over two rounds of PEGylation (Chandradoss et al., 2014). After assembly of a microfluidic flow chamber, slides were incubated for 10 min with T50 buffer (Tris-HCl pH8.0, 50 mM NaCl) containing 5% Tween-20 to further improve slide quality (Pan et al., 2015). Next, the chamber was incubated with 20 μL streptavidin (0.1 mg/ml, Invitrogen, Cat# S-888) for 5 min followed by a washing step with 100 μL of Cas1-Cas2 buffer (50 mM HEPES-NaOH pH7.5, 50 mM KCl, 5 mM MgCl_2_). Biotinylated Cas1-Cas2 were specifically immobilized through biotin-streptavidin linkage by incubating the chamber for 5 min. Remaining unbound biotin-Cas1-Cas2 were flushed away with 100 μL of Cas1-Cas2 imaging buffer (50 mM HEPES-NaOH pH7.5, 50 mM KCl, 5 mM MgCl_2_, glucose oxidase (Sigma, Cat# G2133), 4 mg/ ml Catalase (Roche, Cat#10106810001) and 1mM Trolox ((±)-6-Hydroxy-2,5,7,8-tetramethylchromane-2-carboxylic acid, Sigma, Cat#238813). Immobilized Cas1-Cas2 was incubated with 500 pM labelled DNA at room temperature (23 ± 1 °C) for indicated times.

To visualize the dynamics of PS DNA binding on Cas1-Cas2, Cy3 molecules were excited on an area of 50×50 μm^2^ with a 28% of the full laser power (9 mW) green laser (532nm), while the time resolution was set to 0.1 s. Under these imaging conditions we obtained a high signal-to-noise ratio that allowed us to visualize kinetic intermediates while imaging over time periods of 3.5 min. Under these conditions photobleaching of the donor and acceptor dye during our observation time was minimized.

### Single-Molecule TIRF Data Analysis

A series of CCD images were acquired with laboratory-made software at a time resolution of 0.1 s. Fluorescence time traces were extracted with an algorithm written in IDL (ITT Visual Information Solutions) that picked fluorescence spots above a threshold with a defined Gaussian profile. The extracted time traces were analyzed using custom written MATLAB (MathWorks) and python algorithms. FRET efficiency was defined as the ratio between the acceptor intensity and the sum of the acceptor and donor intensities. To determine the binding rate (koff), the start and end of each binding event were determined (Figure 1C). The start of each event was marked by an abrupt decrease in the donor signal, whereas the end of each event was marked by an abrupt increase in the donor signal (Figure 1C). Selecting the start and end of each event yielded the duration of each event, which was plotted in a histogram. These dwell-time distributions were fitted with a single-exponential decay using maximum-likelihood estimations (Figure S1C). This fit yielded an average dwell-time (Δτ), which was then converted to the binding rate (k_off_ = 1/Δτ). The 95% confidence intervals (errors) of the binding rates were obtained by empirical bootstrap analysis (Dekking, 2005).

The binding frequency was determined by measuring the time from flow-in of the DNA substrate, to the occurrence of the first binding event. These characteristic times were plotted as a cumulative histogram and fitted with a single-exponential decay using maximum-likelihood estimations (Figure S1E and S2D). This fit yielded an average arrival-time (Δτ), which was then converted to the binding rate (k_on_ = 1/Δτ). The 95% confidence intervals (errors) of the binding frequencies were obtained by empirical bootstrap analysis (Dekking, 2005).

To obtain survival rates of the long-lived population, fluorescently labelled PS DNA was incubated for 10 minutes in the microfluidic chamber. After washing the remaining unbound molecules (t=0), the bound population was tracked over time a time of 45 minutes. To avoid photobleaching, short snap-shots of 10 frames were taken over 20 fields of view at each time-point, providing the average number of molecules bound to Cas1-Cas2. For the subsequent analysis, the number of lost molecules at each time-point were subtracted from the total number of molecules bound at t=0. This yielded survival rate curves that were fitted with a single-exponential decay (Figure S1D).

### Quantification and Statistical Analysis

Histograms and fits were generated using OriginPro (OriginLab). The averages and errors representing the number of bound molecules (Figure 1I and S2B), the survival time (Figures 1F and S1D), the accumulative probability of molecule arrival time (Figures 1G, 2A, 2C, 3B, 3E S1E and S2D), dwell time of events (Figures 1H, 2B, 2C, 3C and S1C) and FRET population analysis (Figures 1J, 2D, 5C, S1B, S1H, S2F, S5D and S6E) encompass a minimum of three replicates (n). The errors represent the standard error of the mean (SEM), which was defined as: SEM= σ/√n. The averages and errors displayed in Figures were obtained through bootstrap analysis. In brief, for bootstrap analysis, 104 datasets were generated by random sampling with replacement from the original dataset. Each of these datasets were fitted with the respective fit (indicated in the figure legend) and then used to calculate the average and 95% confidence intervals, which was defined as: CI(.95) = 1.96* σ.

### In Vitro Integration Assays and In Vitro Trimming-Driven Integration Assays

Integration assays with fluorescently labelled PS DNA were performed using 200 nM Cas1-Cas2 complex, 500 nM IHF dimer, 20 nM PS DNA, and 40 nM CRISPR DNA in a final reaction with Cas1-Cas2 integration buffer (50 mM HEPES-NaOH pH7.5, 50 mM KCl, 5 mM MgCl_2_, 5% PEG8000). Cas1-Cas2 complex was incubated with PS DNA to allow complex formation for 30 min at room temperature. Subsequently, IHF dimer was incubated with CRISPR DNA in a separate tube. The reaction was activated by adding the PS-Cas1-Cas2 complex to the IHF-CRISPR DNA mix and incubated for the indicated times at 37°C. For full-site integration experiments of half-site intermediate constructs, 20 nM Half-site integration products were incubated with 500 nM IHF dimer for 30 min at room temperature, followed and by the addition of 200 nM Cas1-Cas2. This mixture was incubated at room temperature for 10 min, allowing Cas1-Cas2 and IHF dimer to assemble with the DNA. Next, 100 nM DNA Pol III, 2.5 U of ExoT or no enzyme was added, followed by incubation at 37°C for indicated times. For trimming-driven integration assays, PS-Cas1-Cas2 and IHF-CRISPR DNA mixtures were combined and incubated at room temperature for 10 min. The combined mixtures were added with or without 100 nM DNA Pol III or 2.5 U of ExoT, followed by incubation at 37°C for indicated times. To quench the reactions, DNA loading buffer (final concentration: 12.5 mM EDTA and 47.5% formamide) was added and thoroughly mixed with samples. The samples were heated at 95°C for 10 min and immediately loaded and run on 15×15 cm^2^–sized 7M Urea denaturing 9% or 12% polyacrylamide 1× Tri-Borate-EDTA (TBE) gels, which was pre-run for 2 h and run for 2-3 h at 370 V in 0.5× TBE buffer. Fluorescence signals from gels were analysed in Amersham™ Typhoon™ biomolecular imager.

### In Vitro Trimming Assays

For in vitro trimming assays, 20 nM PS DNA was incubated with 200 nM Cas1-Cas2 in Cas1-Cas2 integration buffer containing 10 mM DTT and 10% PEG8000 at room temperature for 30 min, and then added with each exonucleases as indicated amounts like below; 1 U of DNA Pol I, 100 nM of DNA Pol III (core) (for holoenzyme, additionally added with 33.3 nM Clamp loader, 200 nM β-clamp and 10 nM DnaBC helicase, but without SSB), 1 U of ExoI, 1 U of RecBCD, 0.5 U of ExoVII or 1 U of ExoT. After incubation at 37°C for indicated times, reaction was quenched with DNA loading buffer and subsequently heated at 95°C for 10 min for 15×15 cm^2^–sized 7M Urea denaturing 20% TBE-PAGE. Fluorescence signals from gels were analysed in Amersham™ Typhoon™ biomolecular imager.

### Electromobility Shift Assays

Binding assays were performed in buffer containing 50 mM HEPES-NaOH pH7.5, 50 mM KCl, 5 mM MgCl2, 5% PEG8000, 5% glycerol, and 1 mM DTT. Each reaction contained 10 nM Dye-labelled PS DNAs or ssDNAs at increasing concentrations (0–200 nM) of Cas1-Cas2. The reactions were incubated at room temperature for 30 min and resolved at 4°C on 4% native agarose gels containing 1× Tris-Acetate-EDTA (TAE) buffer. Fluorescence signals from gels were analysed in Amersham™ Typhoon™ biomolecular imager.

## REFERENCES

Babu, M., Beloglazova, N., Flick, R., Graham, C., Skarina, T., Nocek, B., Gagarinova, A., Pogoutse, O., Brown, G., Binkowski, A., et al. (2011). A dual function of the CRISPR-Cas system in bacterial antivirus immunity and DNA repair. Mol Microbiol 79, 484–502.

Barrangou, R., Fremaux, C., Deveau, H., Richards, M., Boyaval, P., Moineau, S., Romero, D.A., and Horvath, P. (2007). CRISPR provides acquired resistance against viruses in prokaryotes. Science 315, 1709–1712.

Carrico, I.S., Carlson, B.L., and Bertozzi, C.R. (2007). Introducing genetically encoded aldehydes into proteins. Nat Chem Biol 3, 321–322.

Chandradoss, S.D., Haagsma, A.C., Lee, Y.K., Hwang, J.H., Nam, J.M., and Joo, C. (2014). Surface passivation for single-molecule protein studies. J Vis Exp.

Cubbon, A., Ivancic-Bace, I., and Bolt, E.L. (2018). CRISPR-Cas immunity, DNA repair and genome stability. Biosci Rep 38.

Datsenko, K.A., Pougach, K., Tikhonov, A., Wanner, B.L., Severinov, K., and Semenova, E. (2012). Molecular memory of prior infections activates the CRISPR/Cas adaptive bacterial immunity system. Nat Commun 3, 945.

Dekking, M. (2005). A modern introduction to probability and statistics : understanding why and how (London: Springer).

Deveau, H., Garneau, J.E., and Moineau, S. (2010). CRISPR/Cas system and its role in phage-bacteria interactions. Annu Rev Microbiol 64, 475–493.

Diez-Villasenor, C., Guzman, N.M., Almendros, C., Garcia-Martinez, J., and Mojica, F.J. (2013). CRISPR-spacer integration reporter plasmids reveal distinct genuine acquisition specificities among CRISPR-Cas I-E variants of Escherichia coli. RNA Biol 10, 792–802.

Dillard, K.E., Brown, M.W., Johnson, N.V., Xiao, Y., Dolan, A., Hernandez, E., Dahlhauser, S.D., Kim, Y., Myler, L.R., Anslyn, E.V., et al. (2018). Assembly and Translocation of a CRISPR-Cas Primed Acquisition Complex. Cell 175, 934–946 e915.

Drabavicius, G., Sinkunas, T., Silanskas, A., Gasiunas, G., Venclovas, C., and Siksnys, V. (2018). DnaQ exonuclease-like domain of Cas2 promotes spacer integration in a type I-E CRISPR-Cas system. EMBO Rep 19.

Fineran, P.C., and Charpentier, E. (2012). Memory of viral infections by CRISPR-Cas adaptive immune systems: acquisition of new information. Virology 434, 202–209.

Fineran, P.C., Gerritzen, M.J., Suarez-Diez, M., Kunne, T., Boekhorst, J., van Hijum, S.A., Staals, R.H., and Brouns, S.J. (2014). Degenerate target sites mediate rapid primed CRISPR adaptation. Proc Natl Acad Sci U S A 111, E1629–1638.

Garneau, J.E., Dupuis, M.E., Villion, M., Romero, D.A., Barrangou, R., Boyaval, P., Fremaux, C., Horvath, P., Magadan, A.H., and Moineau, S. (2010). The CRISPR/Cas bacterial immune system cleaves bacteriophage and plasmid DNA. Nature 468, 67–71.

Hale, C.R., Zhao, P., Olson, S., Duff, M.O., Graveley, B.R., Wells, L., Terns, R.M., and Terns, M.P. (2009). RNA-guided RNA cleavage by a CRISPR RNA-Cas protein complex. Cell 139, 945–956.

Hamdan, S., Carr, P.D., Brown, S.E., Ollis, D.L., and Dixon, N.E. (2002). Structural basis for proofreading during replication of the Escherichia coli chromosome. Structure 10, 535–546.

Hille, F., Richter, H., Wong, S.P., Bratovic, M., Ressel, S., and Charpentier, E. (2018). The Biology of CRISPR-Cas: Backward and Forward. Cell 172, 1239–1259.

Horvath, P., and Barrangou, R. (2010). CRISPR/Cas, the immune system of bacteria and archaea. Science 327, 167–170.

Hou, Z., and Zhang, Y. (2018). Insights into a Mysterious CRISPR Adaptation Factor, Cas4. Mol Cell 70, 757–758.

Hsiao, Y.Y., Fang, W.H., Lee, C.C., Chen, Y.P., and Yuan, H.S. (2014). Structural insights into DNA repair by RNase T--an exonuclease processing 3’ end of structured DNA in repair pathways. PLoS Biol 12, e1001803.

Huang, Y., Braithwaite, D.K., and Ito, J. (1997). Evolution of dnaQ, the gene encoding the editing 3’ to 5’ exonuclease subunit of DNA polymerase III holoenzyme in Gram-negative bacteria. FEBS Lett 400, 94–98.

Ivancic-Bace, I., Cass, S.D., Wearne, S.J., and Bolt, E.L. (2015). Different genome stability proteins underpin primed and naive adaptation in E. coli CRISPR-Cas immunity. Nucleic Acids Res 43, 10821–10830.

Jackson, S.A., McKenzie, R.E., Fagerlund, R.D., Kieper, S.N., Fineran, P.C., and Brouns, S.J. (2017). CRISPR-Cas: Adapting to change. Science 356.

Kieper, S.N., Almendros, C., Behler, J., McKenzie, R.E., Nobrega, F.L., Haagsma, A.C., Vink, J.N.A., Hess, W.R., and Brouns, S.J.J. (2018). Cas4 Facilitates PAM-Compatible Spacer Selection during CRISPR Adaptation. Cell Rep 22, 3377–3384.

Kim, S., Seo, D., Kim, D., Hong, Y., Chang, H., Baek, D., Kim, V.N., Lee, S., and Ahn, K. (2015). Temporal Landscape of MicroR-NA-Mediated Host-Virus Crosstalk during Productive Human Cytomegalovirus Infection. Cell Host Microbe 17, 838–851.

Kunne, T., Kieper, S.N., Bannenberg, J.W., Vogel, A.I., Miellet, W.R., Klein, M., Depken, M., Suarez-Diez, M., and Brouns, S.J. (2016). Cas3-Derived Target DNA Degradation Fragments Fuel Primed CRISPR Adaptation. Mol Cell 63, 852–864.

Lee, H., Zhou, Y., Taylor, D.W., and Sashital, D.G. (2018). Cas4-Dependent Prespacer Processing Ensures High-Fidelity Programming of CRISPR Arrays. Mol Cell 70, 48–59 e45.

Levy, A., Goren, M.G., Yosef, I., Auster, O., Manor, M., Amitai, G., Edgar, R., Qimron, U., and Sorek, R. (2015). CRISPR adaptation biases explain preference for acquisition of foreign DNA. Nature 520, 505–510.

Loeff, L., Brouns, S.J.J., and Joo, C. (2018). Repetitive DNA Reeling by the Cascade-Cas3 Complex in Nucleotide Unwinding Steps. Mol Cell 70, 385–394 e383.

Lopez-Sanchez, M.J., Sauvage, E., Da Cunha, V., Clermont, D., Ratsima Hariniaina, E., Gonzalez-Zorn, B., Poyart, C., Rosin-ski-Chupin, I., and Glaser, P. (2012). The highly dynamic CRISPR1 system of Streptococcus agalactiae controls the diversity of its mobilome. Mol Microbiol 85, 1057–1071.

Lovett, S.T. (2011). The DNA Exonucleases of Escherichia coli. EcoSal Plus 4.

Marraffini, L.A. (2015). CRISPR-Cas immunity in prokaryotes. Nature 526, 55–61.

Marraffini, L.A., and Sontheimer, E.J. (2008). CRISPR interference limits horizontal gene transfer in staphylococci by targeting DNA. Science 322, 1843–1845.

McGinn, J., and Marraffini, L.A. (2019). Molecular mechanisms of CRISPR-Cas spacer acquisition. Nat Rev Microbiol 17, 7–12.

Moch, C., Fromant, M., Blanquet, S., and Plateau, P. (2017). DNA binding specificities of Escherichia coli Cas1-Cas2 integrase drive its recruitment at the CRISPR locus. Nucleic Acids Res 45, 2714–2723.

Mulepati, S., and Bailey, S. (2013). In vitro reconstitution of an Escherichia coli RNA-guided immune system reveals unidirectional, ATP-dependent degradation of DNA target. J Biol Chem 288, 22184–22192.

Musharova, O., Klimuk, E., Datsenko, K.A., Metlitskaya, A., Logacheva, M., Semenova, E., Severinov, K., and Savitskaya, E. (2017). Spacer-length DNA intermediates are associated with Cas1 in cells undergoing primed CRISPR adaptation. Nucleic Acids Res 45, 3297–3307.

Nunez, J.K., Bai, L., Harrington, L.B., Hinder, T.L., and Doudna, J.A. (2016). CRISPR Immunological Memory Requires a Host Factor for Specificity. Mol Cell 62, 824–833.

Nunez, J.K., Harrington, L.B., Kranzusch, P.J., Engelman, A.N., and Doudna, J.A. (2015a). Foreign DNA capture during CRISPR-Cas adaptive immunity. Nature 527, 535–538.

Nunez, J.K., Kranzusch, P.J., Noeske, J., Wright, A.V., Davies, C.W., and Doudna, J.A. (2014). Cas1-Cas2 complex formation mediates spacer acquisition during CRISPR-Cas adaptive immunity. Nat Struct Mol Biol 21, 528–534.

Nunez, J.K., Lee, A.S., Engelman, A., and Doudna, J.A. (2015b). Integrase-mediated spacer acquisition during CRISPR-Cas adaptive immunity. Nature 519, 193–198.

Pan, H., Xia, Y., Qin, M., Cao, Y., and Wang, W. (2015). A simple procedure to improve the surface passivation for single molecule fluorescence studies. Phys Biol 12, 045006.

Radovcic, M., Killelea, T., Savitskaya, E., Wettstein, L., Bolt, E.L., and Ivancic-Bace, I. (2018). CRISPR-Cas adaptation in Escherichia coli requires RecBCD helicase but not nuclease activity, is independent of homologous recombination, and is antagonized by 5’ ssDNA exonucleases. Nucleic Acids Res 46, 10173–10183.

Redding, S., Sternberg, S.H., Marshall, M., Gibb, B., Bhat, P., Guegler, C.K., Wiedenheft, B., Doudna, J.A., and Greene, E.C. (2015). Surveillance and Processing of Foreign DNA by the Escherichia coli CRISPR-Cas System. Cell 163, 854–865.

Rollie, C., Graham, S., Rouillon, C., and White, M.F. (2018). Prespacer processing and specific integration in a Type I-A CRISPR system. Nucleic Acids Res 46, 1007–1020.

Savitskaya, E., Semenova, E., Dedkov, V., Metlitskaya, A., and Severinov, K. (2013). High-throughput analysis of type I-E CRISPR/ Cas spacer acquisition in E. coli. RNA Biol 10, 716–725.

Schmidt, F., Cherepkova, M.Y., and Platt, R.J. (2018). Transcriptional recording by CRISPR spacer acquisition from RNA. Nature 562, 380–385.

Semenova, E., Savitskaya, E., Musharova, O., Strotskaya, A., Vorontsova, D., Datsenko, K.A., Logacheva, M.D., and Severinov, K. (2016). Highly efficient primed spacer acquisition from targets destroyed by the Escherichia coli type I-E CRISPR-Cas interfering complex. Proc Natl Acad Sci U S A 113, 7626–7631.

Sheth, R.U., and Wang, H.H. (2018). DNA-based memory devices for recording cellular events. Nat Rev Genet 19, 718–732.

Sheth, R.U., Yim, S.S., Wu, F.L., and Wang, H.H. (2017). Multiplex recording of cellular events over time on CRISPR biological tape. Science 358, 1457–1461.

Shiimori, M., Garrett, S.C., Graveley, B.R., and Terns, M.P. (2018). Cas4 Nucleases Define the PAM, Length, and Orientation of DNA Fragments Integrated at CRISPR Loci. Mol Cell 70, 814–824 e816.

Shipman, S.L., Nivala, J., Macklis, J.D., and Church, G.M. (2016). Molecular recordings by directed CRISPR spacer acquisition. Science 353, aaf1175.

Shipman, S.L., Nivala, J., Macklis, J.D., and Church, G.M. (2017). CRISPR-Cas encoding of a digital movie into the genomes of a population of living bacteria. Nature 547, 345–349.

Shmakov, S., Savitskaya, E., Semenova, E., Logacheva, M.D., Datsenko, K.A., and Severinov, K. (2014). Pervasive generation of oppositely oriented spacers during CRISPR adaptation. Nucleic Acids Res 42, 5907–5916.

van der Oost, J., Westra, E.R., Jackson, R.N., and Wiedenheft, B. (2014). Unravelling the structural and mechanistic basis of CRISPR-Cas systems. Nat Rev Microbiol 12, 479–492.

Viswanathan, M., Dower, K.W., and Lovett, S.T. (1998). Identification of a potent DNase activity associated with RNase T of Escherichia coli. J Biol Chem 273, 35126–35131.

Wan, H., Li, J., Chang, S., Lin, S., Tian, Y., Tian, X., Wang, M., and Hu, J. (2019). Probing the Behaviour of Cas1-Cas2 upon Protospacer Binding in CRISPR-Cas Systems using Molecular Dynamics Simulations. Sci Rep 9, 3188.

Wang, J., Li, J., Zhao, H., Sheng, G., Wang, M., Yin, M., and Wang, Y. (2015). Structural and Mechanistic Basis of PAM-Dependent Spacer Acquisition in CRISPR-Cas Systems. Cell 163, 840–853.

Wright, A.V., Liu, J.J., Knott, G.J., Doxzen, K.W., Nogales, E., and Doudna, J.A. (2017). Structures of the CRISPR genome integration complex. Science 357, 1113–1118.

Wright, A.V., Nunez, J.K., and Doudna, J.A. (2016). Biology and Applications of CRISPR Systems: Harnessing Nature’s Toolbox for Genome Engineering. Cell 164, 29–44.

Wu, Y.H., Franden, M.A., Hawker, J.R., Jr., and McHenry, C.S. (1984). Monoclonal antibodies specific for the alpha subunit of the Escherichia coli DNA polymerase III holoenzyme. J Biol Chem 259, 12117–12122.

Xiao, Y., Ng, S., Nam, K.H., and Ke, A. (2017). How type II CRISPR-Cas establish immunity through Cas1-Cas2-mediated spacer integration. Nature 550, 137–141.

Yeeles, J.T., Gwynn, E.J., Webb, M.R., and Dillingham, M.S. (2011a). The AddAB helicase-nuclease catalyses rapid and processive DNA unwinding using a single Superfamily 1A motor domain. Nucleic Acids Res 39, 2271–2285.

Yeeles, J.T., van Aelst, K., Dillingham, M.S., and Moreno-Herrero, F. (2011b). Recombination hotspots and single-stranded DNA binding proteins couple DNA translocation to DNA unwinding by the AddAB helicase-nuclease. Mol Cell 42, 806–816.

Yoganand, K.N., Sivathanu, R., Nimkar, S., and Anand, B. (2017). Asymmetric positioning of Cas1-2 complex and Integration Host Factor induced DNA bending guide the unidirectional homing of protospacer in CRISPR-Cas type I-E system. Nucleic Acids Res 45, 367–381.

Yosef, I., Goren, M.G., and Qimron, U. (2012). Proteins and DNA elements essential for the CRISPR adaptation process in Escherichia coli. Nucleic Acids Res 40, 5569–5576.

Zuo, Y., and Deutscher, M.P. (2002a). Mechanism of action of RNase T. I. Identification of residues required for catalysis, substrate binding, and dimerization. J Biol Chem 277, 50155–50159.

Zuo, Y., and Deutscher, M.P. (2002b). The physiological role of RNase T can be explained by its unusual substrate specificity. J Biol Chem 277, 29654–29661.

