## Supplementary Information Kim & Loeff et al. April 2019 for "Selective Prespacer Processing Ensures Precise CRISPR-Cas Adaptation"

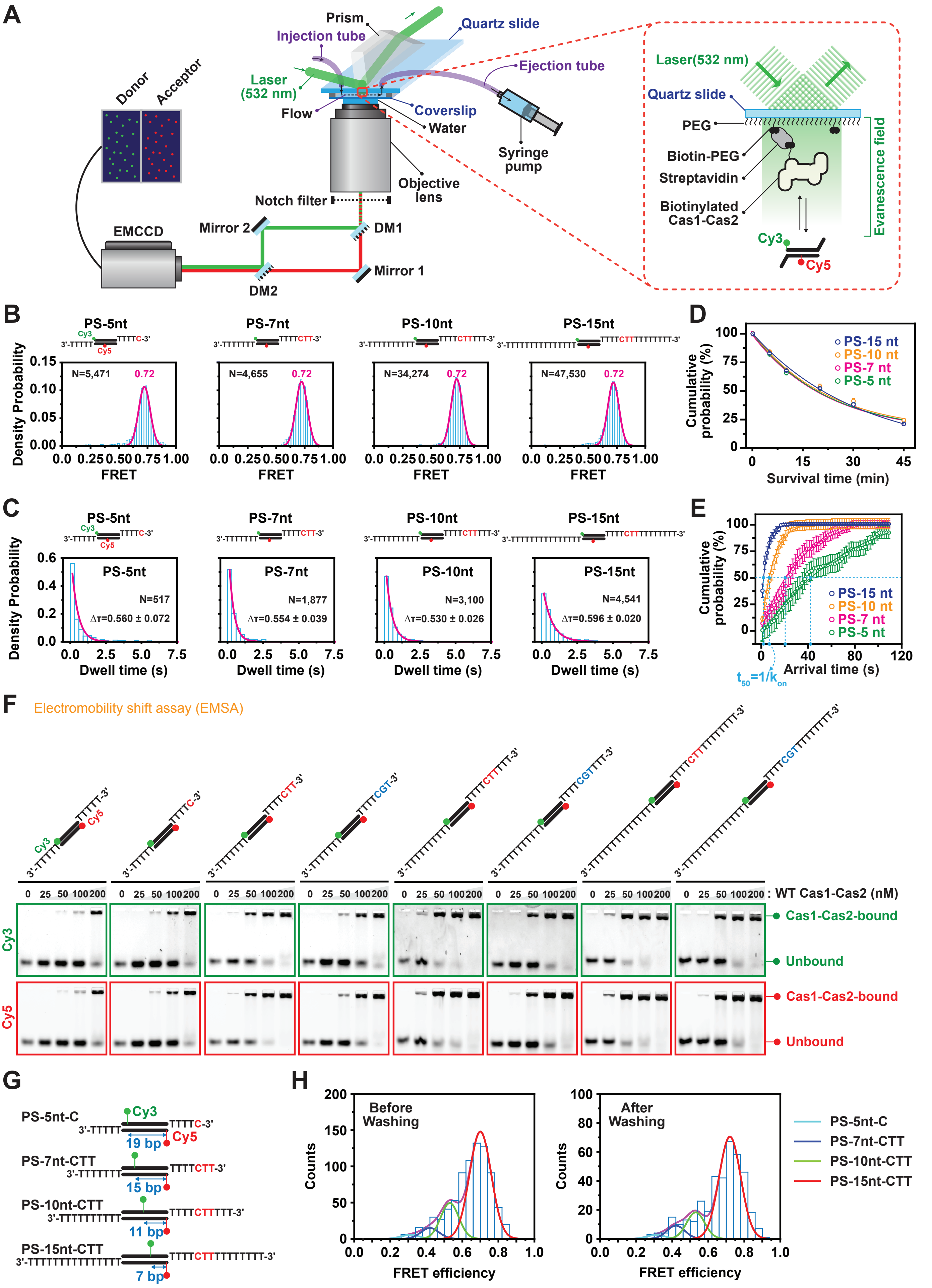
A
Prism
Injection tube
Quartz slide
Laser (532 nm)
Flow
Coverslip
Water
Ejection tube
Syringe pump
Objective lens
Notch filter
Donor
Acceptor
EMCCD
Mirror 2
DM1
Mirror 1
DM2
Laser(532 nm)
Quartz slide
PEG
Biotin-PEG
Streptavidin
Biotinylated Cas1-Cas2
Cy3
Cy5
Evanescence field
B
PS-5nt
PS-7nt
PS-10nt
PS-15nt
N=5,471
N=4,655
N=34,274
N=47,530
0.72
Density Probability
FRET
C
N=517
ΔT=0.560 ± 0.072
N=1,877
ΔT=0.554 ± 0.039
N=3,100
ΔT=0.530 ± 0.026
N=4,541
ΔT=0.596 ± 0.020
Dwell time (s)
D
Cumulative probability (%)
Survival time (min)
PS-15 nt
PS-10 nt
PS-7 nt
PS-5 nt
E
Arrival time (s)
t50=1/kon
F
Electromobility shift assay (EMSA)
0 25 50 100 200 : WT Cas1-Cas2 (nM)
Cy3
Cy5
Cas1-Cas2-bound
Unbound
G
PS-5nt-C
PS-7nt-CTT
PS-10nt-CTT
PS-15nt-CTT
19 bp
15 bp
11 bp
7 bp
H
Before Washing
After Washing
Counts
FRET efficiency
PS-5nt-C
PS-7nt-CTT
PS-10nt-CTT
PS-15nt-CTT

**Figure S1 : Single-molecule analysis of 3'-overhang length-dependent assay and biochemical validation**

**(A)** Experimental set-up of single-molecule TIRF for measuring PS DNA binding kinetics of Cas1-Cas2. **(B)** Histograms for FRET efficiency of selected events observed in 3'-overhang length-dependent smTIRF imaging. **(C)** Dwell-time ($\Delta\tau$) histograms of short-lived single-molecule events of PS DNAs with various 3'-overhang lengths. Related to Figure 1C and 1H. **(D)** Cumulative probability of survival times for various 3'-overhang lengths. The solid lines represent a single-exponential fit. Error bars indicate the 95% confidence interval obtained by bootstrap analysis. Related to Figure 1F. **(E)** Cumulative probability of arrival times of events for various PS 3'-overhang lengths. Solid lines represent single-exponential fit used to determine association binding frequency (kon). Data are represented as mean ± SEM (n = 3). Error bars represent the 95% confidence interval obtained through bootstrap analysis. Related to Figure 1G. **(F)** Electromobility shift assay (EMSA) using various dye-labelled PS DNAs combined with the increasing concentration of wild type Cas1-Cas2. Top strands were labelled with Cy3 at the 5'-ends and bottom strands were labelled with Cy5 at the 5'-ends. Cas1-Cas2-bound and unbound PS DNAs are indicated at the right side of panels. **(G)** Dye-labelling designs of PS DNAs with various 3'-overhang lengths used in the single-molecule competitive Cas1-Cas2 assay. Related to Figure 1J. **(H)** Histograms of the single-molecule competitive Cas1-Cas2 assay using PS DNAs with various 3'-overhang lengths for the comparison of differential populations. Left, before washing including both transient and stable binding events. Right, after washing for measurement of the stably bound population. Each population was fitted with a Gaussian distribution, of which the peak corresponds with the already measured position of the individual constructs. Related to Figure 1J.

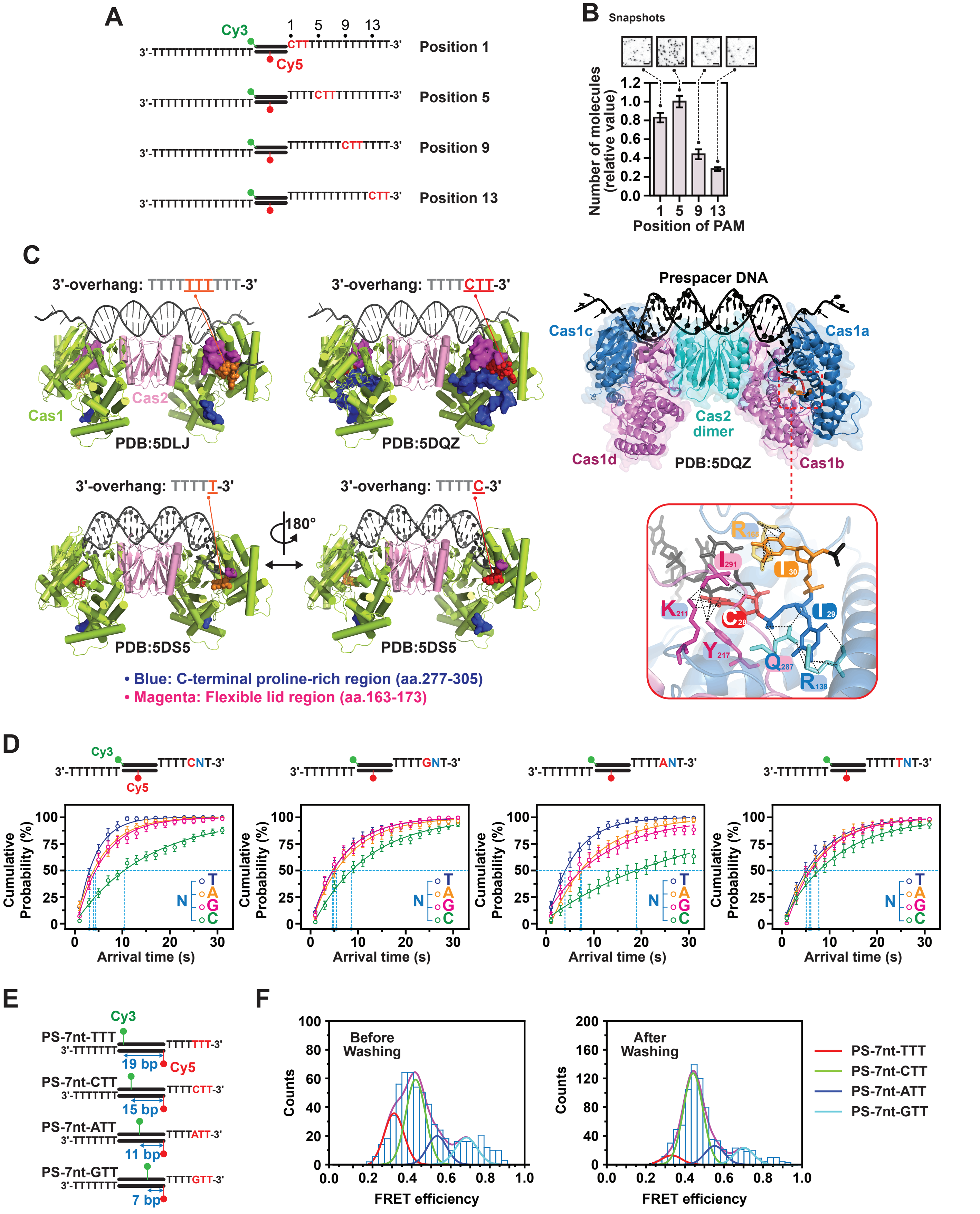

A
Cy3
Cy5
1 5 9 13
3'-TTTTTTTTTTTTTTTT CTTTTTTTTTTTTTT-3'
Position 1
3'-TTTTTTTTTTTTTTTT TTTTCTTTTTTTTTT-3'
Position 5
3'-TTTTTTTTTTTTTTTT TTTTTTTTCTTTTTT-3'
Position 9
3'-TTTTTTTTTTTTTTTT TTTTTTTTTTTTCTT-3'
Position 13
B
Snapshots
Number of molecules (relative value)
Position of PAM
C
3'-overhang: TTTTTTTTTT-3'
PDB:5DLJ
3'-overhang: TTTTCTT-3'
PDB:5DQZ
Cas1
Cas2
Prespacer DNA
Cas1c
Cas1a
Cas2 dimer
Cas1d
Cas1b
PDB:5DQZ
3'-overhang: TTTTT-3'
PDB:5DS5
180°
3'-overhang: TTTTC-3'
PDB:5DS5
• Blue: C-terminal proline-rich region (aa.277-305)
• Magenta: Flexible lid region (aa.163-173)
I291
I30
K211
C28
I29
Y217
Q287
R138
D
3'-TTTTTTT TTTTCNT-3'
3'-TTTTTTT TTTTGNT-3'
3'-TTTTTTT TTTTANT-3'
3'-TTTTTTT TTTTTNT-3'
Cumulative Probability (%)
Arrival time (s)
N T A G C
E
PS-7nt-TTT
3'-TTTTTTT TTTTTTT-3'
19 bp
PS-7nt-CTT
3'-TTTTTTT TTTTCTT-3'
15 bp
PS-7nt-ATT
3'-TTTTTTT TTTTATT-3'
11 bp
PS-7nt-GTT
3'-TTTTTTT TTTTGTT-3'
7 bp
F
Before Washing
After Washing
Counts
FRET efficiency
PS-7nt-TTT
PS-7nt-CTT
PS-7nt-ATT
PS-7nt-GTT

**Figure S2 : Single-molecule analysis and structural insights of PAM-specific PS binding and competitive assays**
**(A)** Designs of Dye-labelling and PAM positions of PS DNAs for various PAM positions in the PS 3'-overhang used in the single-molecule binding assay. **(B)** Quantification of the number of molecules for various PAM positions in the PS 3'-overhang at 30 min after PS DNA addition. Data are represented as mean ± SEM (n = 3). Representative CCD images (acceptor channel) are included as in-sets. **(C)** Structural comparison among Cas1-Cas2 combined with various PS DNAs. PS-Cas1-Cas2 complex with non-PAM (5'-TTT-3', orange) sequence-containing 10-nt-long 3'-overhangs (PDB: 5DLJ, upper left), PAM (5'-CTT-3', red) sequence-containing 7-nt-long 3'-overhangs (PDB: 5DQZ, upper right), and 5-nt-long 3'-overhangs ended by T (PDB: 5DS5, lower left) or C (PDB: 5DS5, lower right) at either 3'-ends are depicted in the right four panels. C-terminal proline-rich tail of Cas1 (blue) and flexible internal lid-like loop region of Cas1 (magenta) are highlighted with indicated colors. PAM recognizing amino acid residues at PAM binding sites in Cas1-Cas2 are zoomed-in at the right panel. C28 (red), T29 (blue) and T30 (orange) in PAM sequence are indicated together with their interacting Cas1 amino acid residues through dotted black lines. **(D)** Cumulative probability of arrival time (s) of events over various sequences at the PAM position in PS 3'-overhang. Solid lines represent single-exponential decay fit used to determine association frequency ($k_{on}$). Data are represented as mean ± SEM (n = 3). Error bars represent the 95% confidence interval obtained through bootstrap analysis. Related to Figure 2A. **(E)** Dye-labelling designs of PS DNAs with various 3'-overhang sequences used in the single-molecule competitive Cas1-Cas2 assay. Related to Figure 2D. **(F)** Histograms of the single-molecule competitive Cas1-Cas2 assay using PS DNAs with various 3'-overhang sequences for the comparison of differential populations. Left, before washing including both transient and stable bindings. Right, after washing for measurement of stable bindings. Each population fits in Gaussian distributions with already measure FRET peaks of each constructs. Related to Figure 2D.

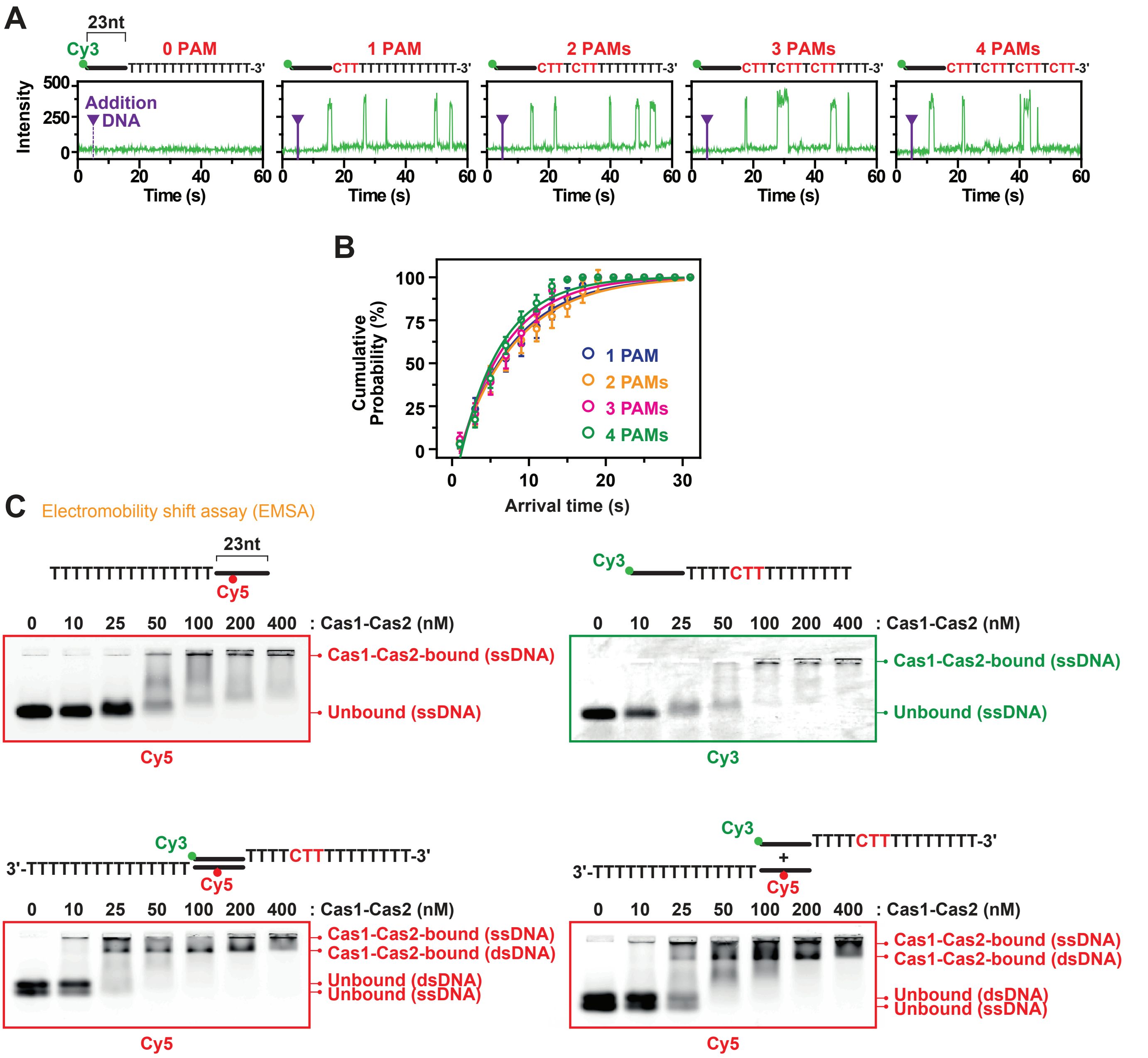


**Figure S3 : Single-molecule trajectories of ssDNA binding of Cas1-Cas2 and biochemical validation**

**(A)** Representative time traces of ssDNA bindings with various number of PAM sites at a single spot in the individual microfluidic chambers. The duration of each binding is measured as the dwell time ($\Delta\tau$) for 60 s. DNA was added at time = 5 s. Related to Figure 3A and 3C. **(B)** Cumulative probability of arrival time (s) of events over the number of PAMs in ssDNAs. Solid lines represent a single-exponential fit used to determine association frequency (kon). Data are represented as mean ± SEM (n = 3). Error bars represent the 95% confidence interval obtained through bootstrap analysis. Related to Figure 3B. **(C)** EMSA using various dye-labelled non-PAM or PAM-containing ssDNAs (upper panels), or already base-paired or sequentially base-paired dsDNAs (lower panels). Cas1-Cas2-bound and unbound ssDNAs or dsDNAs are indicated at the right side of each panel.

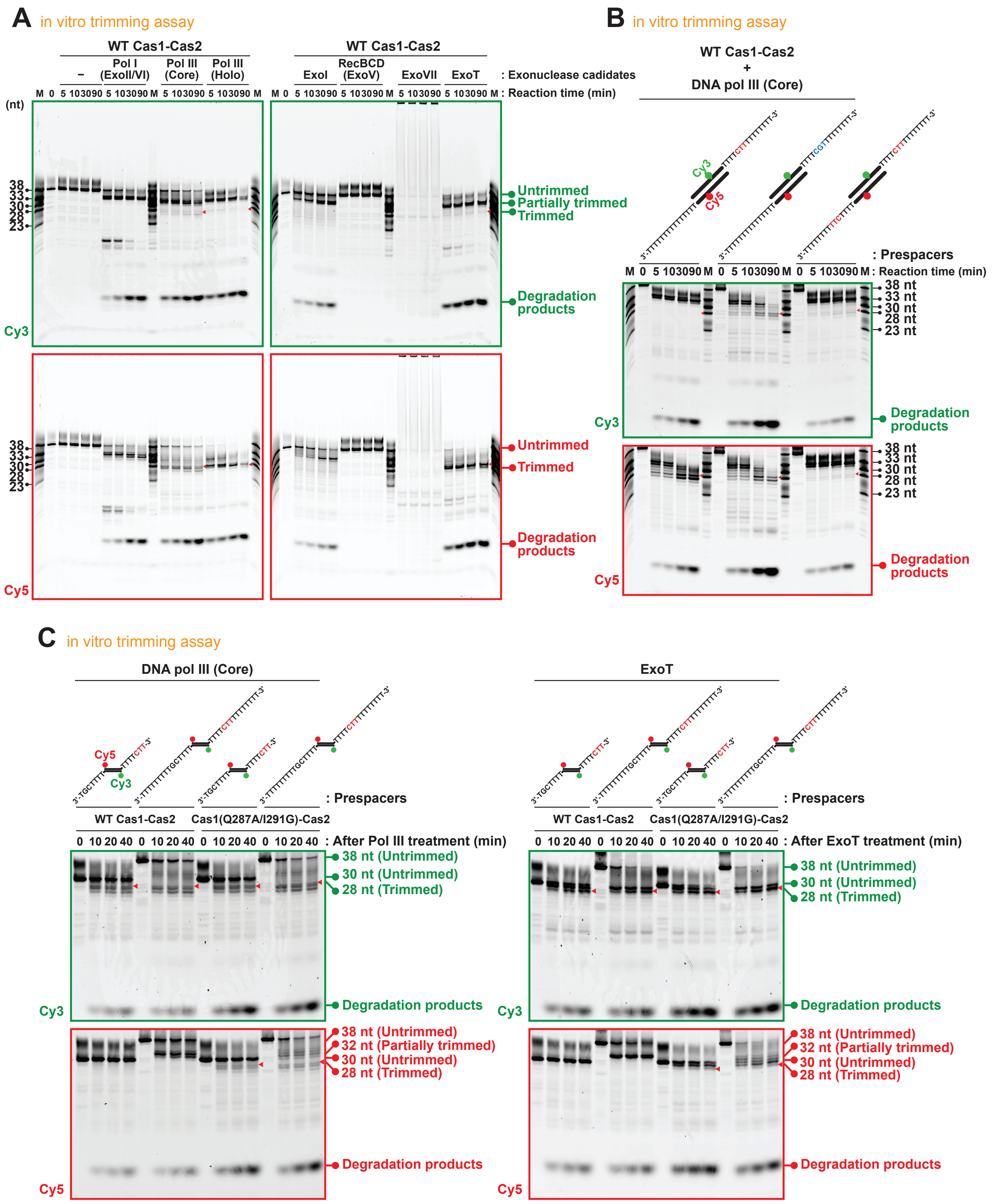


**Figure S4 : In vitro trimming assays**

**(A)** Time-course representation of in vitro trimming assay with various candidate 3'-overhang trimming enzymes. The assay was performed in the same condition as Figure 4B, but samples were collected at 0, 5, 10, 30 and 90 min after reaction with exonucleases. Cy3 and Cy5 signals from 7M Urea denaturing 20% TBE PAGE gels are visualized with Typhoon Scanner and represented as full images. **(B)** In vitro trimming assay with represented PS DNA constructs and DNA Pol III (core) done as (A). **(C)** In vitro trimming assay using represented PS DNA constructs with DNA Pol III (core) (left) or ExoT (right) in the same condition as Figure S4A, but with wild type (WT) or mutant Cas1(Q287A/I291G)-Cas2.

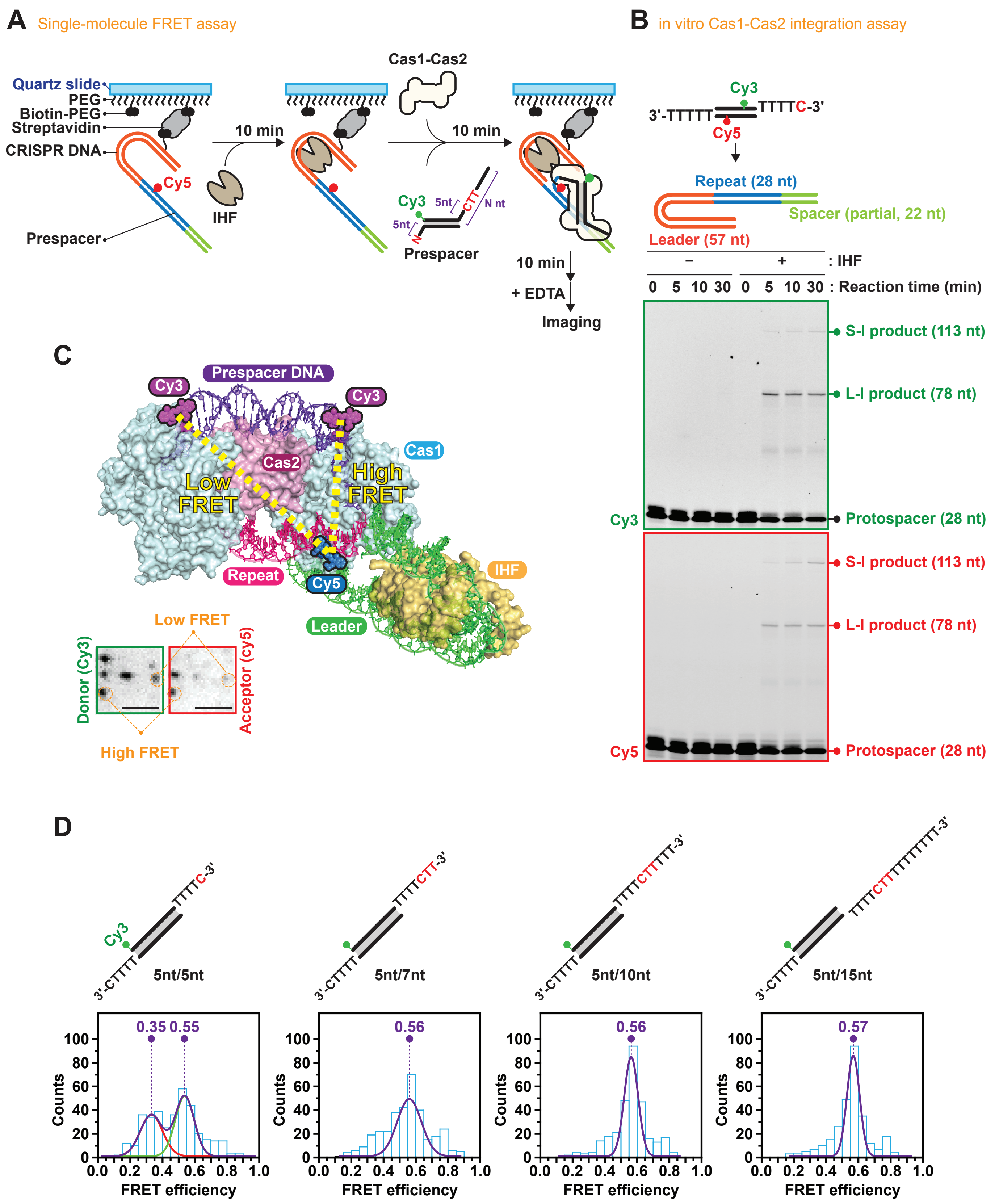


**Figure S5 : In vitro integration assay and smFRET assays**

**(A)** Schematic design of the sample preparation procedure of smFRET assay to observe orientation-sensitive integration. **(B)** In vitro integration assay with or without IHF. Full-site integration in CRISPR DNA (tandemly arrayed 57-nt leader, 28-nt-repeat and 22-nt partial spacer) produces 78-nt leader-side (L-I) products and 113-nt spacer-side (S-I) products by the integration of PS DNA with 28-nt strands at both strands. Cy3 and Cy5 were labelled at the middle of both strands respectively as indicated. **(C)** Expected FRET values for the orientation-sensitive smFRET-based assay. Structural file (PDB: 5WFE) was used for the modeling. Representative CCD images in donor (in green box) and acceptor (in red box) channels are included as in-sets and indicated with representative high and low FRET values. **(D)** Histograms of FRET-selected events, which is related to Figure 5C. Gaussian fitting was applied to extract high and low FRET populations as differential ratios.

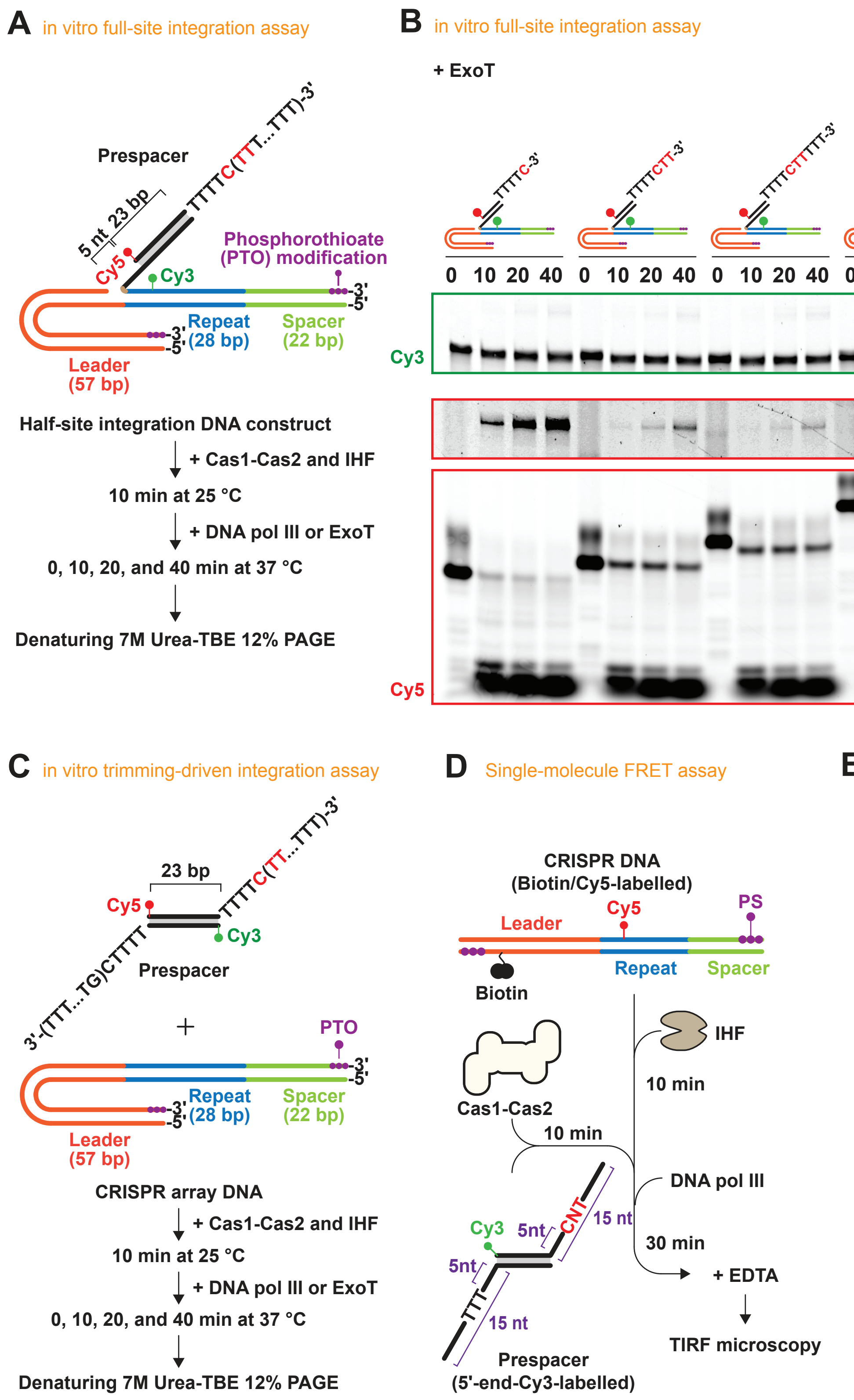


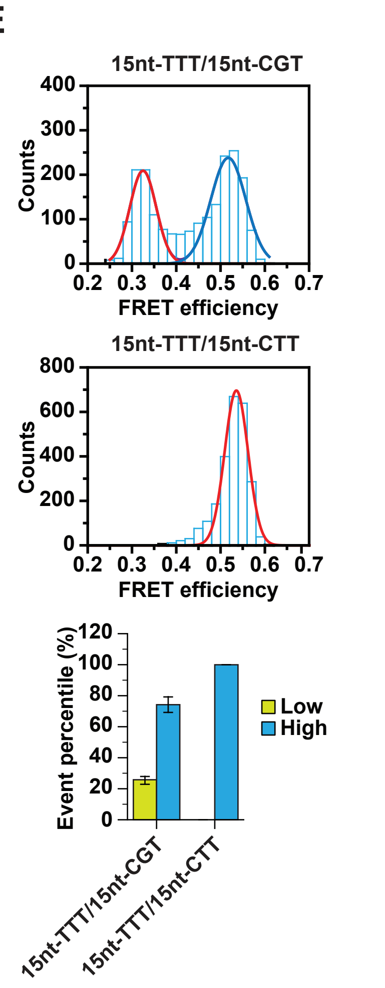


**Figure S6 : In vitro integration assay and smFRET assay**
**(A)** Experimental procedure of in vitro full-site integration assay. Half-site intermediate products are depicted. Various top strands of PS DNAs are represented in Figure 6A. Phosphorothioate (PTO) modification was used in the last three backbone phosphodiester bonds from 3'-ends of CRISPR array DNA for preventing from 3'-5' exonuclease degradation. Half-site intermediate products used in each condition are presented in Figure 6A. **(B)** Gel images of the in vitro trimming-driven full-site integration assay with ExoT. **(C)** Experimental procedure of in vitro trimming-driven integration assay. Designs of PTO-modified CRISPR array DNA and dye-labelled PS DNAs are depicted. PS DNAs used in each condition are represented also in Figure 6B. **(D)** Experimental procedure of trimming-driven integration smFRET assay. **(E)** Histograms of high and low FRET populations are represented and analysed by Gaussian fitting. Bar graphs are shown to compare different proportions of each integration orientation between non-PAM and PAM-containing PS DNAs. Data are represented as mean ± SEM (n = 6).

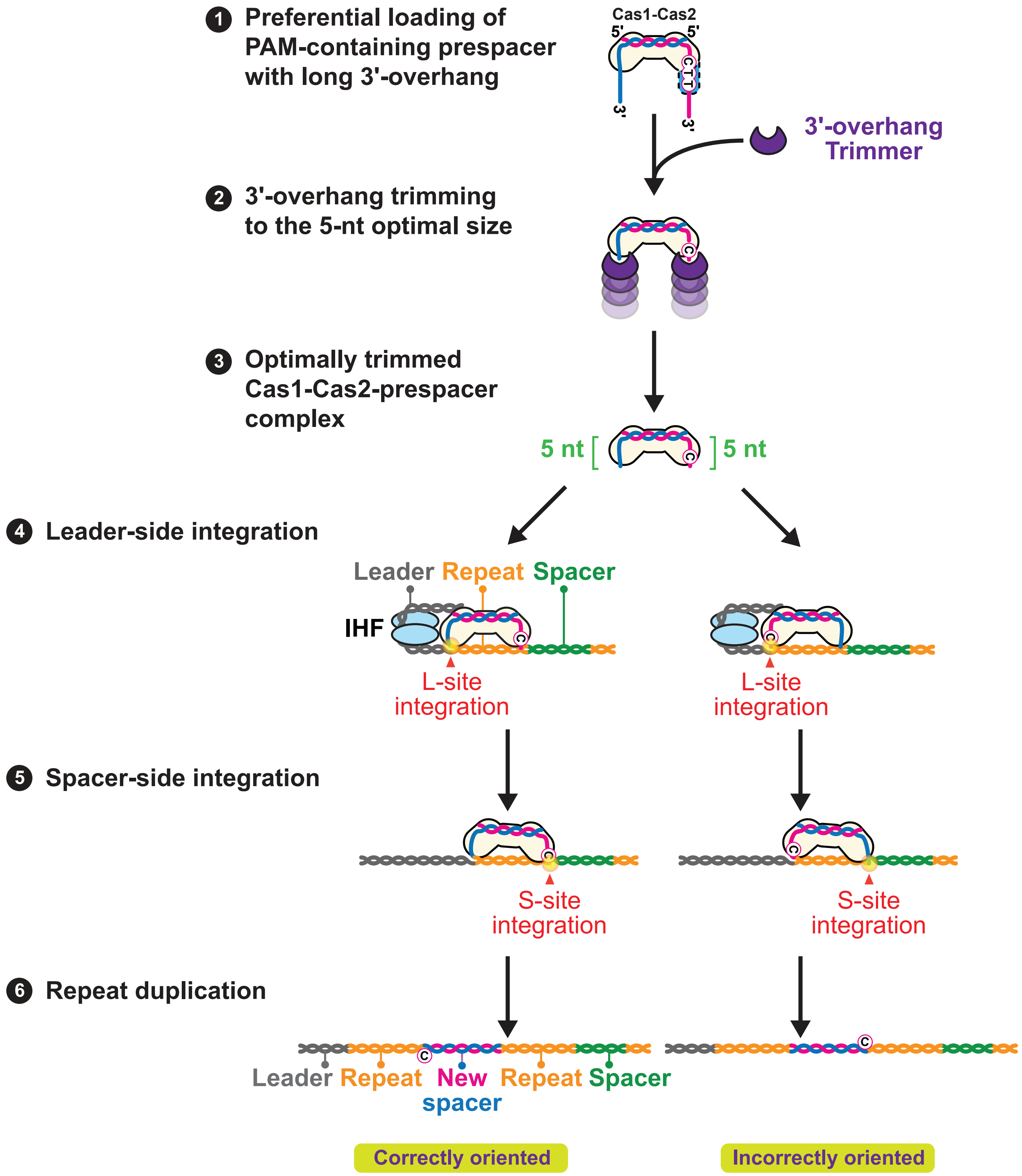


**Figure S7 : Conventional unbiased trimming model of spacer acquisition**
In this model, 3'-overhangs of PAM-containing Cas1-Cas2-selected PS precursors are trimmed symmetrically into 5-nt canonical length. PAM-derived 3'-end of trimmed PS can be integrated in either leader-side or spacer-side integration sites eventually. Possibility of correct orientation in the newly integrated spacers is theoretically 50%, which doesn't fit into physiologically relevant results.
